# Efficient sampling designs to assess biodiversity spatial autocorrelation : should we go fractal ?

**DOI:** 10.1101/2022.07.29.501974

**Authors:** F. Laroche

## Abstract

Quantifying the autocorrelation range of species distribution in space is necessary for applied ecological questions, like implementing protected area networks or monitoring programs. However, the power of spatial sampling designs to estimate this range is negatively related with other objectives such as estimating environmental effects acting upon species distribution. Mixing random sampling points and systematic grid (‘hybrid’ designs) is a classic solution to make a trade-off. However, fractal designs (i.e. self-similar designs with well-identified scales) could make an even better compromise, because they cover a wide array of possible autocorrelation range values across scales. Using maximum likelihood estimation in an optimal design of experiments approach, we compared errors of hybrid and fractal designs when simultaneously estimating an effect acting upon a response variable and the residual autocorrelation range. We found that Pareto-optimal sampling strategies depended on the feasible grid mesh size (FGMS) over the study area, given the sampling budget. When the FMGS was shorter than expected autocorrelation range values, grid design was the best option on all criteria. When the FMGS was around or larger than expected autocorrelation range values, the choice of designs depended on the effect under study. Fractal designs outperformed hybrid designs when studying the effect of a monotonic environmental gradient across space, while grid design was more efficient for other types of question. Beyond the niche identified in our analysis, fractal designs may also appear interesting when studying response variables with more heterogeneous spatial structure across scales, and when considering more practical criteria of performance such as the distance needed to cover the design.

## Introduction

Autocorrelation has a double status in the study of biodiversity patterns (Legendre, 1993). On the one hand, it is often seen as a nuisance, generating biases in regression models that seek to link covariates to spatial patterns of biodiversity (Lennon, 2000). Many techniques to control these undesirable effects are available, and now well popularized among ecologists (Dormann et al., 2007). On the other hand, spatial autocorrelation may also be viewed as the signature of some endogeneous process driving biodiversity patterns (McGill, 2010). In particular, it is often interpreted through the prism of limited dispersal. For instance, auto-regressive modelling of species occupancy in metapopulation ecology (ter Braak et al., 1998; Bardos et al., 2015; Prugh, 2009; Ranius et al., 2010) or isolation by distance patterns on neutral markers in population genetics (Ouborg et al., 1999; Vekemans and Hardy, 2004; Manel and Holderegger, 2013) are often used to draw conclusion on species colonization or dispersal abilities. This interpretation of autocorrelation range can be further reinforced in multi-taxonomic studies, when estimated values are found to be correlated to species dispersal traits (Bonada et al., 2012). From this perspective, the accurate assessment of autocorrelation range has important implications to assess the functional connectivity of habitat networks (Tischendorf and Fahrig, 2000) or build efficient biodiversity monitoring strategies (Rhodes and Jonzén, 2011).

Virtual ecology (Zurell et al., 2010) offers a way to test whether sampling designs can accurately detect or quantify effects of interest, before implementing it in the field. Beyond the question of assessing the power of empirical designs, a virtual ecology analysis contributes to clearly formulating the set of questions associated to a design. Many virtual studies have focused on evaluating the potential of various sampling strategies to estimate the mean population density of a species (e.g. Perret et al. (2022)), the total abundance or the total species richness of a taxon (e.g. Thiele et al. (2023)) within a sampling area. However, only few virtual studies focused on efficient designs to accurately estimate the autocorrelation range of biodiversity variables. An early example is a study by Ferrandino (2004), which analyzed how various sampling schemes could quantify the aggregation of a pathogen distribution among host plants. The main result was that a regular grid sampling design could better estimate the total incidence of the pathogen over the studied area, while more irregular designs (random or fractal) with the same number of points could yield better estimates of the pathogen aggregation, for a wider range of true cluster sizes. Fractals showed the best performance on the latter aspect. Similarly, Bijleveld et al. (2012) found that a grid design was the best choice to estimate spatial or temporal trends on the mean of a target field of values while random design was better at estimating autocorrelation parameters. The authors further showed that a hybrid strategy, mixing randomly chosen sites with a grid, stood as a Pareto-optimal solution on the trade-off between the conflicting objectives (i.e. changing to other designs necessarily generated performance loss on either objective). However, they did not include fractal designs in their analysis. From a species community perspective, Marsh and Ewers (2013) suggested that fractal sampling designs could be an interesting option to study the autocorrelation range of species composition among communities through diversity partitioning or distance-decay patterns (Lande, 1996; Nekola and White, 1999)). They found that fractal designs lead to non-parametric estimates of distance decay-curves more similar to an intensive control than other classic strategies (regular grid and random design), calling for further investigation.

Fractal designs are characterized by a self-similar property (Mandelbrot, 1983; Falconer, 2003): sub-parts of the design look like a contraction of the total design (see Fig. 2 below). Thanks to this property, a single fractal design can cover discrete, contrasted spatial scales with a relatively low sampling effort. They may offer a practical way to study autocorrelation over a broad set of possible ranges. Ewers et al. (2011) emphasized the importance of such feature: it should contribute to create versatile long-term designs, suitable for many resarch questions covering taxa with contrasted dispersal abilities, home range sizes and sensitivity to environment, and facilitate the study of links between scales. Given this high potential, we deemed necessary to compare fractal designs with hybrid designs (including pure grid and pure random designs) regarding their ability to simultaneouly estimate autocorrelation range and effects acting on a target variable in space. By doing so, we hoped to synthesize the pros and cons associated to fractal sampling strategy in terms of estimation power, and identify when then may constitute a valuable alternative compared to more classic strategies on that respect.

We used the statistical framework of optimal design of experiments (Müller et al., 2012), which has been repeatedly used to build and compare designs of temporal (Archaux and Bergès, 2008) or spatio-temporal (see Hooten et al. (2009) and references therein) biodiversity surveys. This approach has rarely been applied to the specific problem of quantifying spatial autocorrelation though. An exception is the study by Müller (2007), which focused on the problem of detecting autocorrelation with a test using Moran index (Moran, 1950), but did not consider the estimation error associated to the corresponding range. Here, we focused on estimation errors associated to an effect acting upon a response variable and to the residual autocorrelation range respectively. We used maximum likelihood estimation, which offers a powerful heuristic to theoretically explore the estimation accuracy of sampling designs through the analysis of the inverse Fisher matrix (Abt and Welch, 1998). Zhu and Stein (2005) used this approach to numerically search for best sampling positions to recover autocorrelation parameters of a random field. They found that emerging designs with lowest error on estimates of autocorrelation parameters differed from random design, and tended to conciliate aggregated points at the center of the surveyed area with points scattered close to the frontier. Such designs might be viewed as harbouring dicrete scales, hence reinforcing the interest of explicitly assessing fractal designs’ performance.

When comparing hybrid and fractal designs, we had two expectations based on previous litterature: (i) hybrid designs should consitute a continuous set of intermediary Pareto-optimal designs between grid and random designs, meaning that when the proportion of random points increases from 0 (grid design) to 1 (random design), the accuracy of the mean estimate of the random field should decrease while the accuracy of the autocorrelation range estimate should increase; (ii) fractal designs should be better than other designs at estimating small autocorrelation ranges when they are built to harbour contrasted scales, hence creating new Pareto-optimal solution focused on autocorrelation range estimation.

## Methods

All the methods described below have been implemented using the *R* language (R Core Team, 2023). Code with detailed comments and examples is available on an online, public repository (doi: 10.5281/zenodo.10245301). Some visualizations of the response variable presented there used the package *mvtnorm* (Genz and Bretz, 2009).

### Spatial sampling designs

All spatial sampling designs harboured *N* = 27 sampling points. Sampling points were spread within an area of study shaped as an equilateral triangle with a side length of 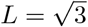 distance units (Figs. 1, 2).

**Figure 1.**
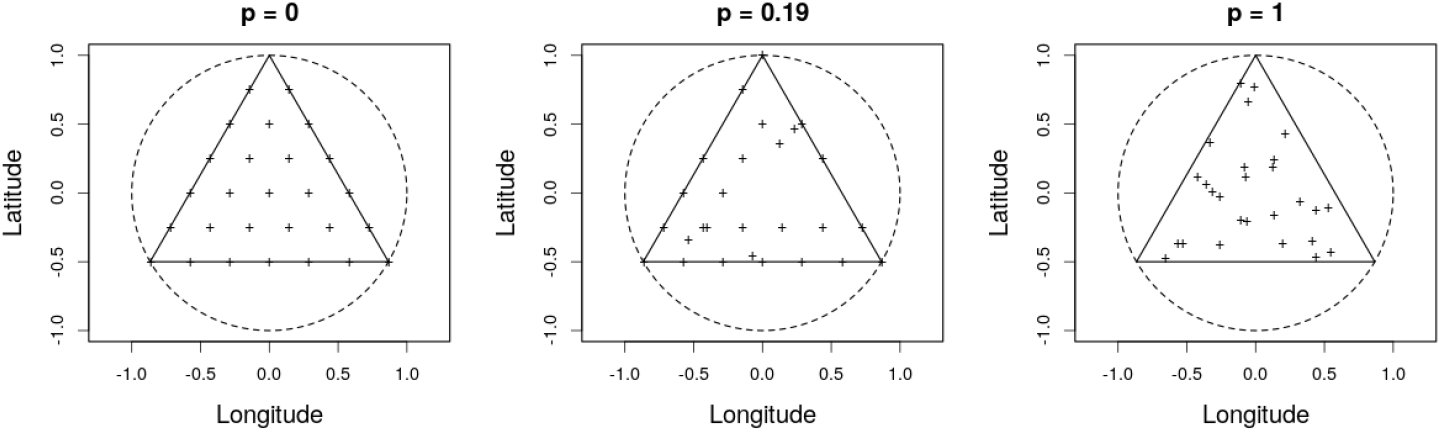
Effect of the proportion of random sites *p* on hybrid sampling designs. The triangle is the area of study. Crosses show the position of the 27 sampling points. The middle panel corresponds to *p* = 5/27 ≈ 0.19.

**Figure 2.**
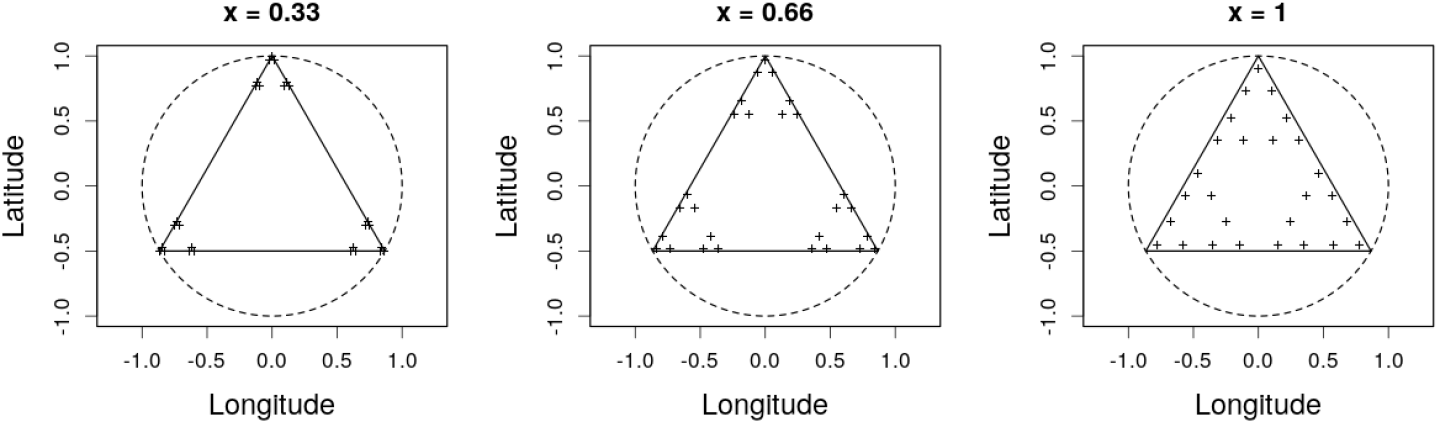
Effect of the contraction parameter *x* on fractal sampling designs. The triangle is the area of study. Crosses show the position of the 27 sampling points.

Grid designs were obtained by generating a triangular grid matching the area of study with mesh size equal to *L/*6 distance units (the ‘feasible grid mesh size’ below), hence obtaining 28 sampling points. Then one point was removed at random to obtain *N* = 27. Hybrid designs were defined by choosing a fraction *p* of sites at random within a grid design and resampling their new position at random in the area of study. Here we consider the *N* + 1 = 28 values of *p ∈* {0, 1*/N*, 2*/N*, …, 1}. Note that *p* = 1 yields a pure random design (Fig. 1). Since all hybrid designs included a random components, we considered 30 replicates for each value of *p*, which led to 28 × 30 = 840 hybrid designs in total.

Following Marsh and Ewers (2013), we simulated fractal designs using an iterated function systems (Falconer, 2003) based on three similarities of the complex plane: 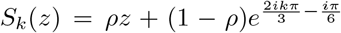 for *k* ∈ {0, 1, 2}. A sampling design was obtained by iterating three times the system starting from a seed at the center of the area, hence yielding a sampling design with *N* = 3^3^ = 27 plots. We varied the parameter *ρ* across designs. The parameter *ρ* drives the ratio between the size of a part of the design and the size of the larger, auto-similar set of plots it belongs to. The values of *ρ* considered in the study were :

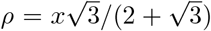 with 28 distinct *x* values evenly spaced on a log-scale from *x* = 10^−1.5^ to *x* = 1. We called *x* the ‘contraction parameter’ of fractal design below. Note that *x* > 1 would generate a sampling design with overlapping sub-components, which we considered as an undesirable property. The largest value *x* = 1 yields a sampling design that is a subsample of a regular grid with mesh size of c.a. *L/*10. Lower *x* yielded more irregular fractal sampling designs (Fig. 2).

Most of the analyses presented in next sections focus on the statistical power of fractal and hybrid sampling designs to estimate some parameters driving the response of a variable of interest. However, statistical designs also differ on more practical aspects such as the effort needed to cover all the sampling sites. We computed the minimum distance necessary to cover all the sampling sites of a design as a proxy of its practicability. This single proxy does not cover all the dimensions that make a design practicable — and covering all these dimensions would go much beyond the scope of the present work — but computing this simple criterion can still reveal potential trade-offs between practical and statistical advantages of various designs.

Comparing the minimum distance needed to cover grid and fractal designs will prove of particular importance below. The minimum distance for grid design is (ArticleS1 supplementary information, section 4):

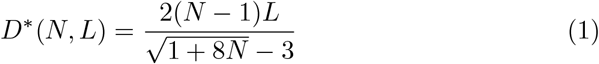

where *N* is the number of sampling points in the grid and *L* is the side length of sampling area. Note that although hybrid designs harbour 27 sampling sites in previous sections, the underpinning grid has *N* = 28 sampling sites.

When *ρ* < 1/3, which is equivalent to *x* < 0.72, the shortest spanning path length of a fractal design is :

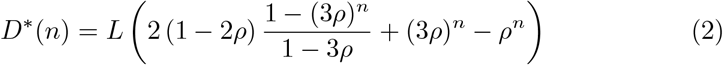

where *n* is the number of iteration to generate the design (*n* = 3 in previous sections). For *x* > 0.72, we kept this distance as a conservative estimate of the minimum distance needed to cover the fractal, keeping in mind that the true minimum distance could be even smaller.

### Modeling the observed variable

We assumed that observations of interest were driven by an environmental gradient with auto-correlated residual variation. The vector of observations at each sampling points ***Y*** = (*Y*_1_, …, *Y*_*N*_) followed the model :

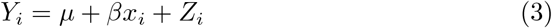

where *μ, β* are parameters in ℝ, *x*_*i*_ is an environmental covariate and ***Z*** = (*Z*_1_, …, *Z*_*N*_) is taken from a Gaussian random field with mean 0 and harbouring an exponential variogram (Cressie, 1993) without nugget effect. This meant that the covariance between *Z*_*i*_ and *Z*_*j*_ was:

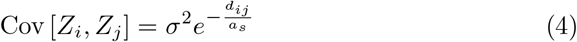

where *σ, a*_*s*_ *∈* ℝ^+*∗*^, *d*_*ij*_ is the distance between sampling points *i* and *j*. Parameter *a*_*s*_ corresponded to the autocorrelation range. We considered 30 distinct *a*_*s*_ values evenly spaced on a log-scale between 10^−2^ and 10^2^.

### Modeling the environmental covariate

We modelled the environmental covariate using a product of two sine waves along the latitudinal and longitudinal axes, generating a checkerboard pattern. We varied the period and the phase of the checkerboard to create five contrasted situations in our analyses below (Fig. 3).

**Figure 3.**
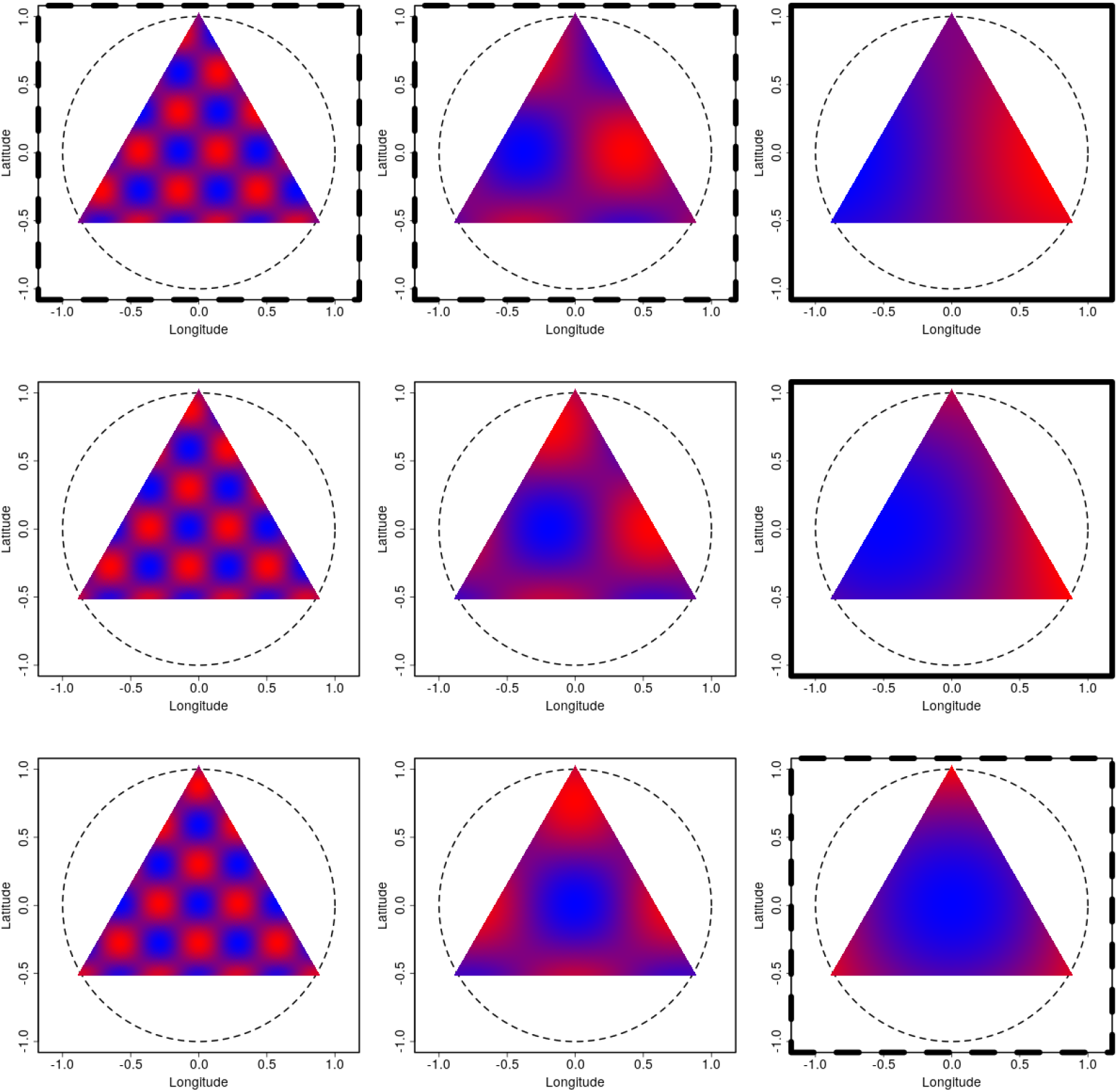
Various spatial profiles for the environmental variable generated with a bidimensional checkerboard. The darkest blue color corresponds to an environmental value of 0, the brightest red color corresponds to an environmental value of 1. The period of the checkerboard increases from left to right, the shift to the right increases from up to bottom. The environmental profiles that are presented in results are highlighted with a thick frame. Those with a continuous frame correspond to monotonic profiles in space, while dashed frames those with a dashed frame correspond to markedly non-monotonic profiles in space.

### Maximum likelihood estimation

We considered two distinct estimation problems of ecological interest : (i) estimating the mean (*μ*) and the autocorrelation range (*a*_*s*_) of the response variable, in the absence of environmental effect (*β* = 0); (ii) estimating the effect of the environmental covariate (*β*) on the response variable and the residual autocorrelation range (*a*_*s*_).

#### Problem 1 : mean versus autocorrelation range

Here we assumed that that the observed covariate had a constant mean in space, following the simplified version of model (3):

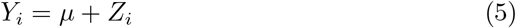

The aim was to accurately estimateg the range of spatial aurocorrelation *a*_*s*_ and the intercept *μ*. The statistical model used to estimate parameters was model (5), i.e. we did not consider potential errors on model specification. When comparing estimation errors on intercept and autocorrelation range, we considered an exponentially-transformed intercept *ν* = *e*^*μ*^ to obtain the same domain of definition ℝ^+*∗*^ as *a*_*s*_. Summarizing parameters in a vector ***θ*** = (*ν, σ, a*_*s*_), we focused on the maximum likelihood estimate 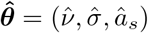.

#### Problem 2 : environmental effect versus autocorrelation range

Here the observed covariate followed model (3). The aim was to accurately estimate the range of spatial aurocorrelation *a*_*s*_ and the slope of the environmental effect *β*. The statistical model used to estimate parameters was a reformulation of model (3) where we assumed that the environmental covariate has been centered in the dataset :

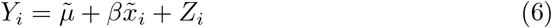

where 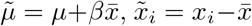 and 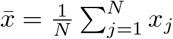. We did not consider potential errors on model specification. When comparing estimation errors on intercept and autocorrelation range, we considered an exponentially-transformed slope *γ* = *e*^*β*^ to obtain the same domain of definition ℝ^+*∗*^ as *a*_*s*_. Summarizing parameters of this model in a vector 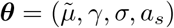, we focused on the maximum likelihood estimate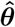.

### Estimation errors

Estimation error on a parameter *θ* (possibly *ν, γ, a*_*s*_ depending on the problem) is quantified through the relative root mean square error:

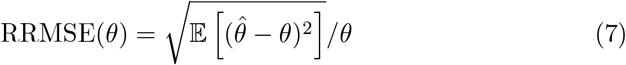

In the context of stationary Gaussian random fields without nugget, it is known that the diagonal terms of *ℐ* (***θ***)^*−*1^, where *ℐ* (***θ***) is the Fisher information matrix of the model with true parameters ***θ***, yield a qualitatively good approximation of the quadratic error on parameters in ***θ***. By ‘qualitatively’, we mean that it allows to correctly rank designs according to their accuracy, even for moderate sample sizes (Abt and Welch, 1998; Zhu and Stein, 2005). We therefore used the diagonal terms of *ℐ* (***θ***)^*−*1^ as a theoretical approximation of quadratic error of 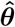 below. The error on parameter *θ*_*i*_ was thus approached by RRMSE 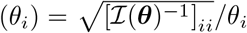. For hybrid designs, errors were averaged across replicates to obtain a single error value per parameter of the model of interest for each value of *p*.

### Comparing designs using Pareto fronts

When comparing designs on their ability to accurately estimate several parameters at a time, we relied on the concept Pareto-optimality, defined as follows. Let generically denote the parameters of interest ***θ*** = (*θ*_1_, *θ*_2_). Denote RMSE_*S*_(*θ*_*i*_) the error of sampling design *S* at estimating parameter *θ*_*i*_ (where *i* = 1, 2). Generally, a design *S*_0_ is said to be Pareto-optimal within the set of designs 𝒮 at estimating the parameters in ***θ*** if and only if there is no other design in 𝒮 that would yield a lower estimation error on all the terms of ***θ***. Formally, Pareto optimality of *S*_0_ is thus expressed as :

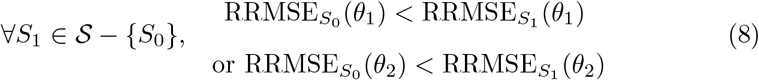

Because we considered designs that could show levels of errors numerically close, we modified this definition to include an idea of ‘noticeable’ difference in estimation error. In our study, a design *S*_0_ is said to be Pareto-optimal within the set of designs 𝒮 if and only if:

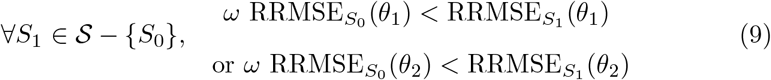

where *ω* = 0.99 is a 1% tolerance factor on errors.

For each pattern of environmental covariate and each value of autocorrelation range *a*_*s*_ we searched for designs that were Pareto-optimal within the set of designs of their own type (fractal or hybrid respectively), and those that were Pareto optimal wihtin the set of all the designs considered in the study.

For each pattern of environmental covariate, we also searched for designs that were Pareto-optimal when considering an ‘average’ level of error across a range of *a*_*s*_ values. As detailed in results, the value *a*_*s*_ = *L/*6 was associated to a marked transition in the qualitative patterns of design errors. We thus considered a range of *a*_*s*_ values centered on this value on a log-scale, starting from 0.1 × *L*/6 up to 10 × *L*/6. Because the level of RRMSEs strongly fluctuates in magnitude across the range of *a*_*s*_ values, directly computing a mean of errors across the range of *a*_*s*_ values is unappropriate. We resorted to first converting RRMSEs of designs at a given *a*_*s*_ value into ranks, and then averaging out the ranks attained by a design at each *a*_*s*_ value to obtain an averaged error rank on each parameter. Ultimately, the average error ranks were used in an analysis of Pareto-optimality.

## Results

### Computing predicted errors

**Problem 1** — The relative root mean squared error associated to 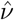 and 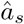 derived from the Fisher information matrix (see Article S1 in Supporting Information, section 1) were:

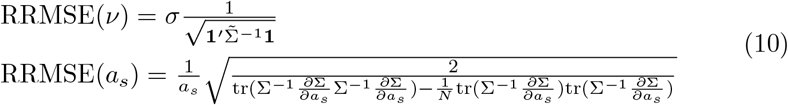

RRMSEs did not depend on *ν*, hence we set *ν* = 1 without loss of generality. RRMSE(*a*_*s*_) did not depend on *σ*, while RRMSE(*ν*) was proportional to *σ*. Because we were interested in the ranking of designs, which remains identical up to a multiplicative constant, we set *σ* = 1 without loss of generality in our conclusions either.

**Problem 2** — The relative root mean squared error associated to 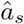 and 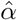 derived from the Fisher information matrix (see Article S1 in Supporting Information, section 1) were:

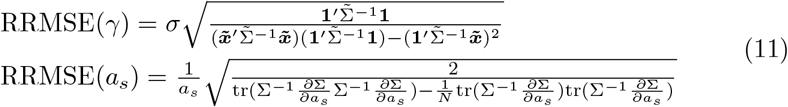

RRMSEs did not depend on 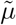 and *γ*, hence we set *μ* = 0 and *γ* = 1 throughout the study without loss of generality. RRMSE(*a*_*s*_) did not depend on *σ* while RRMSE(*γ*) was proportional to *σ*. Here again, because we were interested in the ranks of designs, which are identical up to a multiplicative constant, we set *σ* = 1 without loss of generality in our conclusions either.

Note that RRMSE(*a*_*s*_) is identical in problems 1 and 2 (eqs. (10) and (11)).

### Theoretical analysis of asymptotic errors

We performed a theoretical analysis of errors when *a*_*s*_ took extreme values (small or large). We found that RRMSE(*a*_*s*_) increased towards +*∞* as *a*_*s*_ became small, irrespective of considered design (see Article S1 section 2 in Supporting Information). However, the speed of increase varied across designs. Denoting *d*_min_ the smallest distance among two distinct sampling points, designs with smaller *d*_min_ yielded smaller RRMSE(*a*_*s*_) at low *a*_*s*_ values. The grid design (hybrid design with *p* = 0) maximized *d*_min_ (see Article S1 in Supporting Information, section 3) and was thus expected to yield consistently higher RRMSE(*a*_*s*_) than other designs as *a*_*s*_ *→* 0. Fractal designs could harbour arbitrarily small *d*_min_ by decreasing contraction parameter *x*, and were thus expected to reach lower RRMSE(*a*_*s*_) than any hybrid designs when *x* is sufficiently small.

In problem 1, RRMSE(*ν*) converged towards 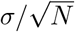 as *a*_*s*_ became small, irrespective of the sampling design. In our case, *σ* = 1 and *N* = 27, which yielded RRMSE(*ν*) ≈ 0.19. Given our tolerance factor on Pareto optimality, we thus expected all designs to become Pareto-optimal for problem 1 as *a*_*s*_ → 0. In problem 2, RRMSE(*γ*) converged towards 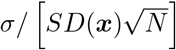 where *SD*(***x***) is the standard deviation of the environmental covariate ***x*** across sampling points. Sampling designs that would maximize the variance in ***x*** while harbouring small values of *d*_min_ should therefore be Pareto-optimal.

For very large *a*_*s*_ values, RRMSE(*a*_*s*_) converged towards 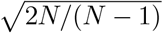 (≈ 1.44 when *N* = 27), irrespective of the sampling design (see Article S1 in Supporting Information, section 2). In problem 1, RRMSE(*ν*) converged to *σ* (= 1 in our example), irrespective of the sampling design. In problem 2, RRMSE(*γ*) converged to 0, irrespective of the sampling design. In either problem, all the sampling designs were thus expected to converge towards very similar performance as *a*_*s*_ increased, and thus be a Pareto-optimal strategy. Regarding RRMSE(*ν*), this phenomenon of convergence among designs had already been shown on simulations in a previous study (Perret et al., 2022).

### Numerical analysis of Pareto fronts in problem 1

We discussed our results according to two main situations below : (i) *a*_*s*_ smaller than the feasible grid mesh size (i.e. *a*_*s*_ < *L*/6 ≈ 0.29); (ii) *a*_*s*_ larger than the feasible grid mesh size (*a*_*s*_ > *L*/6).

#### Fractal designs alone

When *a*_*s*_ was below the feasible grid mesh size, RRMSE(*a*_*s*_) of fractal designs showed a U-shaped pattern as *x* increased while RRMSE(*ν*) decreased as *x* increased (*a*_*s*_ = 0.09 in Fig. 4). Therefore, only an upper range of *x* values corresponded to Pareto-optimal strategies within fractal designs (*a*_*s*_ = 0.09 in Fig. 4; Fig. 5). This upper range of Pareto-optimal *x* values shrinked as *a*_*s*_ increased towards *L/*6 (Figs. 4, 5). As *a*_*s*_ increased above *L/*6, only an intermediary range of *x* led to Pareto-optimal sampling strategies within fractal designs. This range gradually broadened with *a*_*s*_ until the point where all designs became Pareto-optimal (Fig. 5), as theoretically expected.

**Figure 4.**
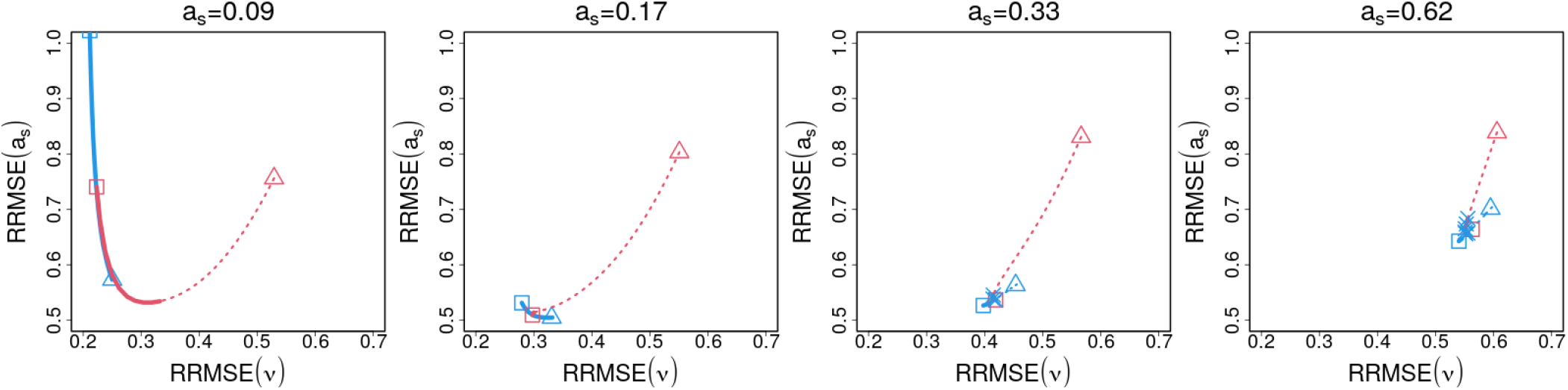
Pareto fronts of fractal and hybrid sampling designs at selected levels of autocorrelation range *a*_*s*_ in problem 1. Lines show the RRMSEs of fractal (red line, tracking the 28 values of *x*) and hybrid (blue line, tracking the 28 values of *p*) designs. The end of lines materialized with a triangle corresponds to the most irregular design (*x* = 10^−1.5^ for fractal designs; *p* = 1 for hybrid designs). The end of lines materialized with a square corresponds to the most regular design (*x* = 1 for fractal designs; *p* = 0 for hybrid designs). Dashed parts of lines (and open end symbols) show designs that are not Pareto-optimal within their own type, while solid parts of lines (and solid end symbols) show those that are. Blue crosses show fractal designs that were Pareto-optimal within their own type and are not anymore when introducing hybrid designs.

**Figure 5.**
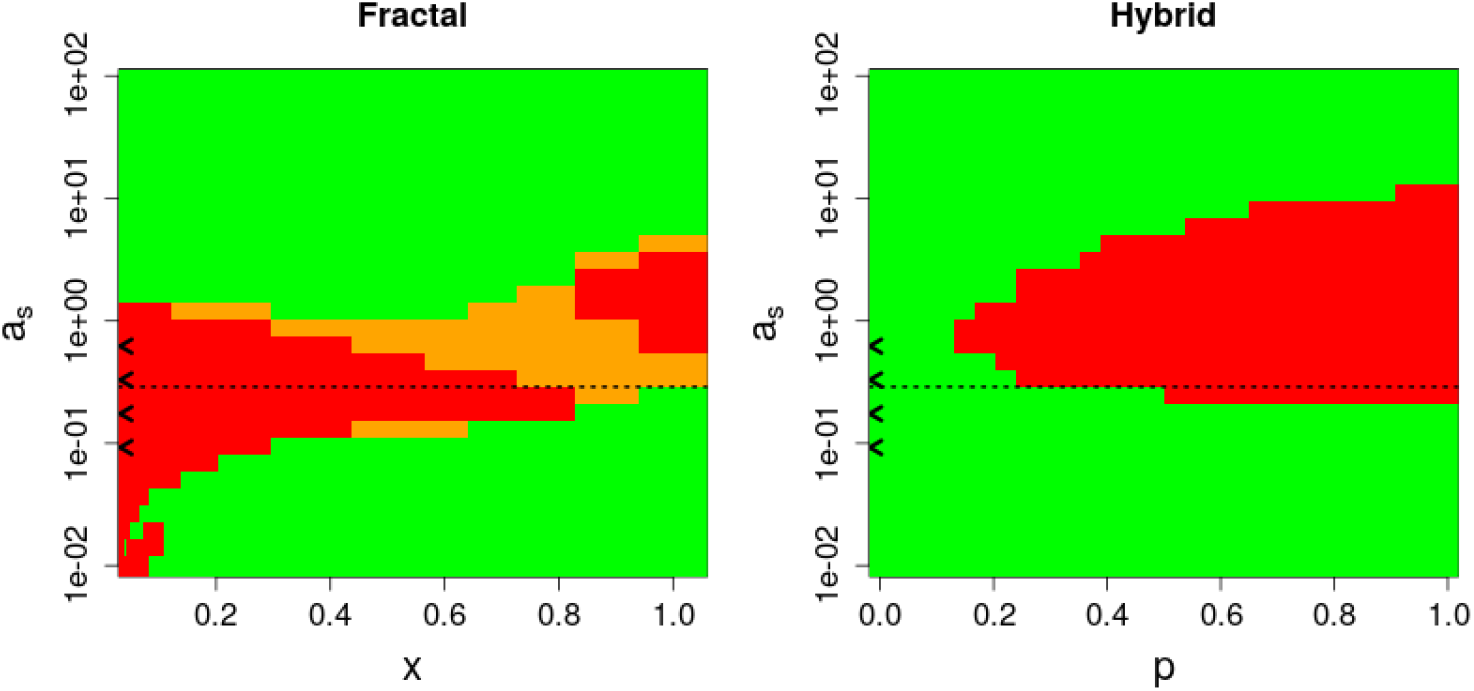
Pareto fronts of sampling designs as a function of autocorrelation range *a*_*s*_ in problem 1. In the left panel, colored pixels indicates, for given *a*_*s*_ and *x* values, whether the corresponding fractal sampling design is Pareto-optimal within fractal and hybrid sampling designs altogether (green pixels) or within fractal sampling designs only (orange pixels). Red pixels show fractal designs that are not Pareto optimal within the set of fractal sampling strategies. Similarly, in the right panel, colored pixels indicates, for given *a*_*s*_ and *p* values, whether the corresponding hybrid sampling design is Pareto-optimal within fractal and hybrid sampling designs altogether (green pixels) or within hybrid sampling designs only (orange pixels). Red pixels show hybrid designs that are not Pareto optimal within the set of hybrid sampling strategies. Therefore, in either panel, green and orange pixels show designs that are Pareto-optimal within their own type, and orange pixels shows the fraction of these designs that are eliminated when introducing the other type of design in the comparison. The horizontal dashed line *a*_*s*_ = *L/*6 is the size of feasible grid mesh size, it shows the limit between two situations discussed in main text. The *a*_*s*_ values illustrated in Figure 4 are reported on the ordinate axis using chevrons.

#### Hybrid designs alone

When *a*_*s*_ was below the feasible grid mesh size, RRMSE(*a*_*s*_) of hybrid designs decreased while RRMSE(*ν*) increased as *p* increased (e.g. *a*_*s*_ = 0.09, 0.17 in Fig. 4). Therefore, any value of *p* yielded a Pareto-optimal strategy within hybrid designs (Fig. 5). As *a*_*s*_ increased above *L/*6, the variation of RRMSE(*a*_*s*_) rapidly shifted from decreasing to increasing with *p* while the profile of RRMSE(*ν*) remained unchanged. Therefore, only the most regular designs remained Pareto-optimal among hybrid designs (*a*_*s*_ = 0.33, 0.62 in Fig. 4 and Fig. 5). When *a*_*s*_ further increased, all the hybrid designs gradually came back to the Pareto front, like any other designs, as theoretically expected (Fig. 5).

#### Fractal and hybrid designs together

When *a*_*s*_ was below the feasible grid mesh size, fractal designs never excluded a hybrid design (Fig. 5, right panel) and, reciprocally, hybrid designs only rarely excluded a fractal designs (Fig. 5, left panel). The coexistence of fractal and hybrid designs on the global Pareto front stemmed from the fact that fractal designs either lead to larger RRMSE(*ν*) and lower RRMSE(*a*_*s*_) than hybrid designs, or generated errors comparable to hybrid designs on both parameters (see e.g. *a*_*s*_ = 0.09 in Fig. 4). By contrast, as *a*_*s*_ values increased above grid mesh size, hybrid designs led to the exclusion of fractal designs from the Pareto-front as long as the errors of designs still had opportunity to noticeably vary (*a*_*s*_ = 0.33, 0.62 in Fig. 4; Fig. 5, left panel).

### Numerical analysis of Pareto fronts in problem 2

The five environmental patterns (Fig. 3) could actually be split in two groups with homogeneous error patterns : one group included the two patterns where the environmental covariate had a monotonic profile (continuous frames in Fig. 3) and one group included the three patterns where the environmental covariate had a markedly non-monotonic profile (dashed frames in Fig. 3). We presented our results using this grouping (monotonic versus non-monotonic) in the text, while Fig. 6 shows results for each individual environmental pattern.

**Figure 6.**
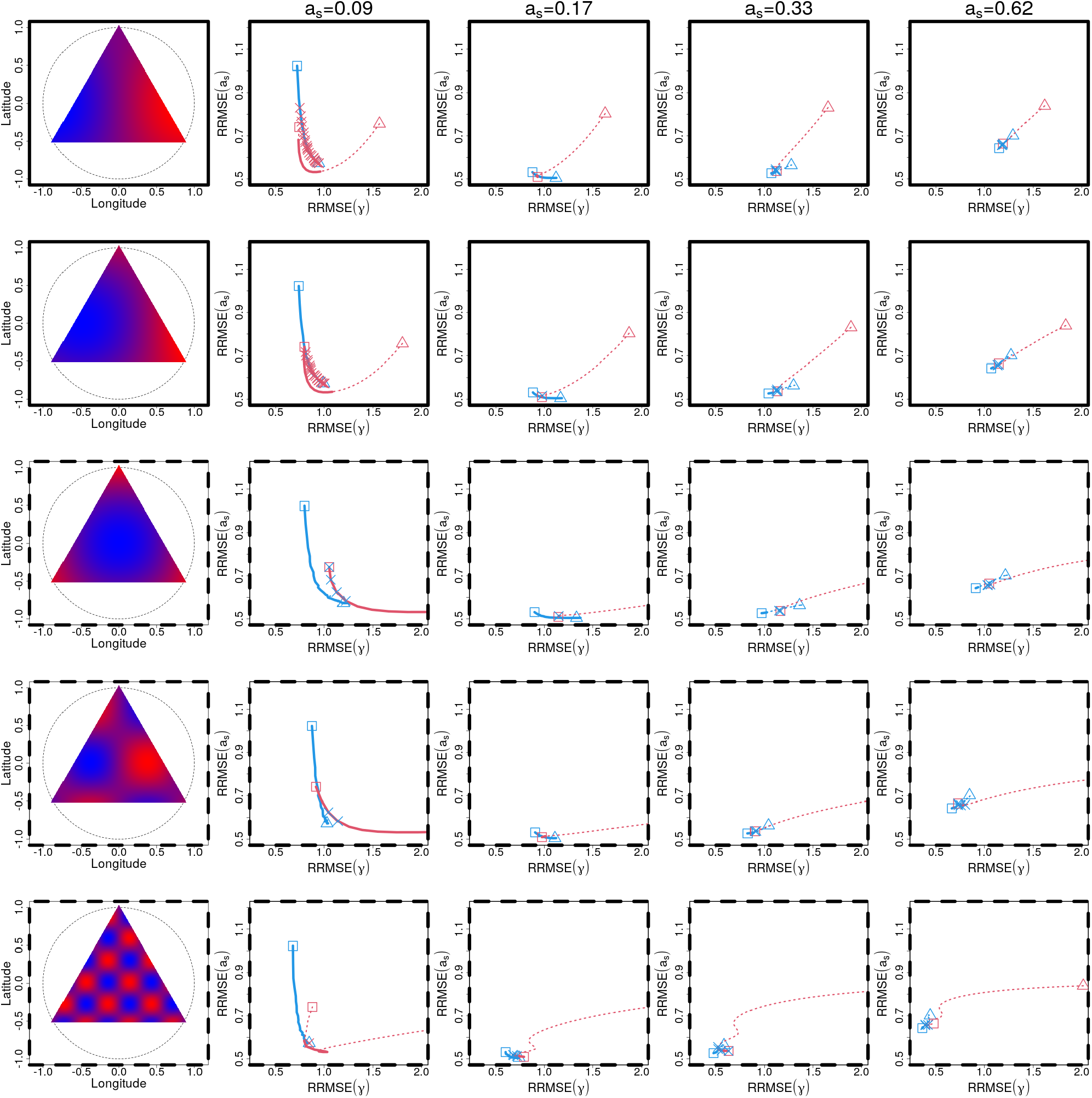
Pareto front of fractal and hybrid sampling designs for various levels of autocorrelation range and environment profiles. The first column show the environment profile considered in the row. Rows with continuous (respectively dashed) frames correspond to monotonic (respectively non-monotonic) profiles in space. Columns two to four show Pareto fronts for increasing levels of autocorrelation range (same legend as in Figure 4). The red triangle (irregular end of fractal designs) can be outside the panel to the right. Red crosses show hybrid designs that are Pareto-optimal within their own type but are not when introducing fractal designs.

### Monotonic environment

#### Fractal designs alone

When *a*_*s*_ was smaller than the feasible grid mesh size, RRMSE(*γ*) harboured a U-shaped pattern as *x* increased (slightly visible for *a*_*s*_ = 0.09 in the two first lines of Fig. 6). Because RRMSE(*a*_*s*_) also harboured a U-shaped pattern, only fractal sampling designs with intermediary *x* comprised between the minima for RRMSE(*a*_*s*_) and RRMSE(*γ*) were Pareto optimal within the set of fractal designs and this range of Pareto-optimal *x* values gradually shifted towards higher values as *a*_*s*_ increased (Fig. 7). When *a*_*s*_ increased above the feasible grid mesh size, we observed an upper range of Pareto-optimal *x* values which broadened as *a*_*s*_ increased until encompassing all fractal designs, as their estimation errors all converged towards the same value (Fig. 7).

**Figure 7.**
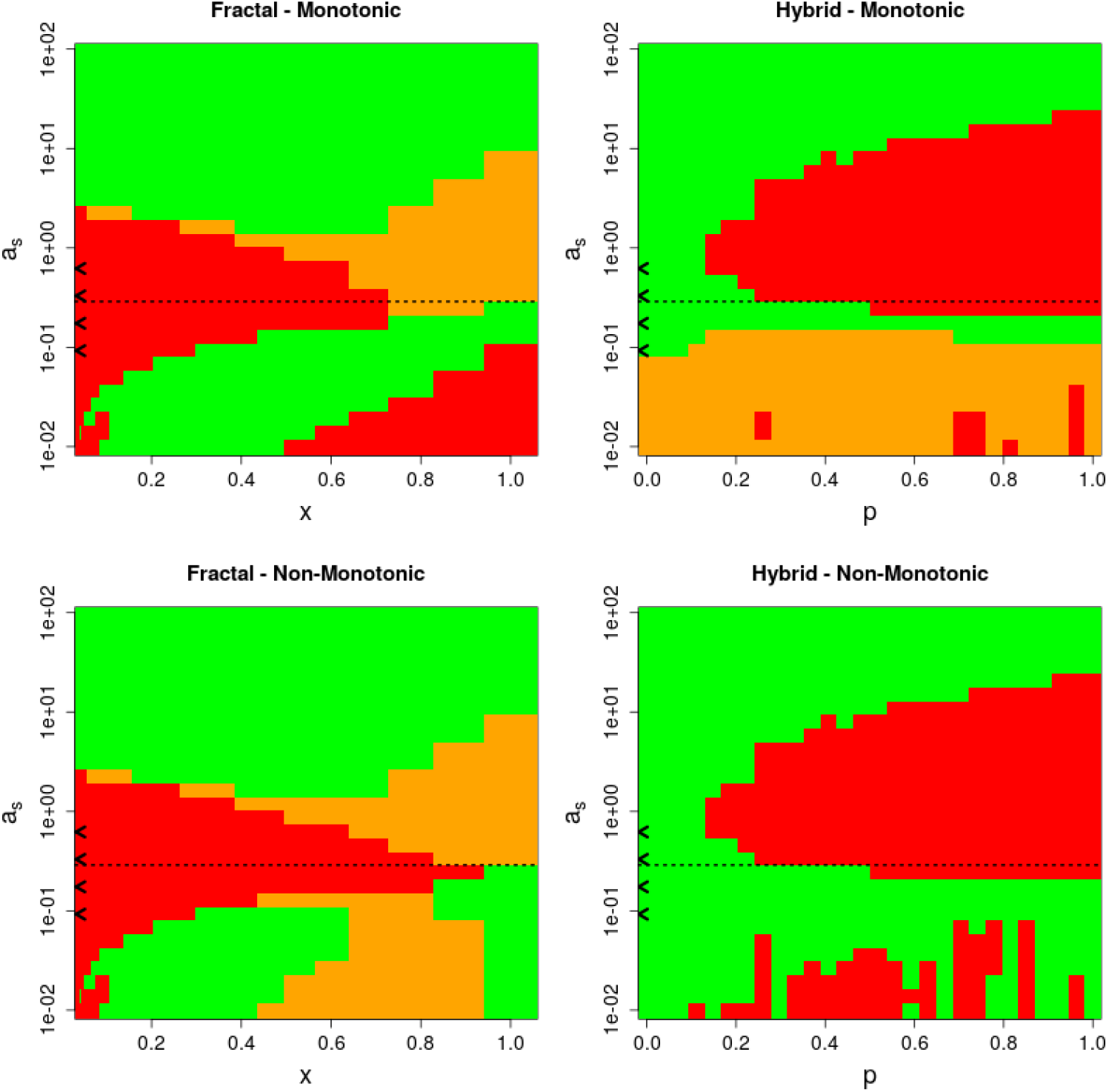
Pareto fronts of sampling designs in problem 2. Panels correspond to one type of design (fractal or hybrid) in one type of environment (monotonic or non-monotonic). We selected first line of Fig. 6 as a representative of monotonic environments, and the fourth line as a representative of non-monotonic environments. In each panel, the horizontal axis presents the regularity parameter of the design (*x* or *p* for fractal or hybrid designs respectively). Colored pixels indicates, for given *a*_*s*_ and regularity parameter values, whether the design is Pareto-optimal within fractal and hybrid designs altogether (green pixels) or within its own type only (orange pixels). Red pixels show designs that are not Pareto optimal within their own type. Therefore, green and orange pixels show designs that are Pareto-optimal within their own type, and orange pixels shows the fraction of these designs that are eliminated when introducing the other type of design in the comparison. The horizontal dashed line *a*_*s*_ = *L/*6 is the size of grid deign mesh size, it shows the limit between two situations discussed in main text.

#### Hybrid designs alone

When *a*_*s*_ was smaller than the feasible grid mesh size, RRMSE(*γ*) monotonically increased with *p*, while RRMSE(*a*_*s*_) decreased. Therefore, any hybrid design along the gradient of *p* was a Paretooptimal strategy (*a*_*s*_ = 0.09, 0.17 in the first two lines of Fig. 6; Fig. 7). When *a*_*s*_ increased above the feasible grid mesh size, the RRMSE(*γ*) remained monotonically increasing with *p* (*a*_*s*_ = 0.33, 0.62 in the first two lines of Fig. 6), while we saw before that RRMSE(*a*_*s*_) abruptly shifted from decreasing to increasing. Therefore, only the most regular designs (lower range of *p*) remained on the Pareto front (Fig. 7). This range broadened when *a*_*s*_ further increased because the estimation errors of all hybrid designs converged towards the same value.

#### Fractal and hybrid designs together

When *a*_*s*_ was smaller than the feasible grid mesh size, fractal designs excluded a large fraction of hybrid designs from the Pareto front, especially those with a high proportion of random points (first two lines of *a*_*s*_ = 0.09 in Fig. 6; Fig. 7, upper right panel). By contrast, when *a*_*s*_ increased above the feasible grid mesh size, hybrid designs with low *p* led to the exclusion of fractal designs from the Pareto-front for all *a*_*s*_ values where the errors of designs still had opportunity to noticeably vary (first two lines of *a*_*s*_ = 0.33, 0.62 in Fig. 6; Fig. 7, left panel).

### Non-monotonic environment

#### Fractal designs alone

When *a*_*s*_ was smaller than grid mesh size, RRMSE(*γ*) harboured a globally decreasing pattern as *x* increased (rows three to five in Fig. 6), although variation could be more irregular at high *x* values in environment with highest ruggedness (row five in Fig. 6). Because RRMSE(*a*_*s*_) harboured a U-shaped profile, the upper range of *x* values formed a range of Pareto optimal strategy within fractal designs in environments with moderate ruggedness (e.g. *a*_*s*_ = 0.09 in rows three and four of Fig. 6; Fig. 7). This upper range of *x* values shrinked as *a*_*s*_ increased towards grid mesh size *L/*6. When *a*_*s*_ increased above *L/*6, the pattern of optimal *x* values was qualitatively similar to the monotonic environment case (Fig. 7).

#### Hybrid designs alone

Like for the case of monotonic environment depicted above, RRMSE(*γ*) monotonically increased with *p* irrespective of the value of *a*_*s*_ (Fig. 6). Consequently, the situation was here again qualitatively similar to the monotonic environment case.

#### Fractal and hybrid designs together

Fractal designs rarely excluded hybrid designs from the Pareto front (Fig. 7, bottom right panel), except in environment with high ruggedness where grid design with high degree of randomness could occasionally be excluded (*a*_*s*_ = 0.09 in row five of Fig. 6). When *a*_*s*_ was smaller than grid mesh size, hybrid designs excluded a large intermediary fraction of fractal designs from the Pareto front (sometimes all of them), mainly because fractal design showed markedly higher errors at estimating the slope *γ* (Fig. 7). Fractal design with lower *x* could persist when they harboured error very similar to some hybrid designs while fractal designs with highest *x* score could persist because they ourperformed hybrid designs at estimating autocorrelation range *a*_*s*_ (e.g. *a*_*s*_ = 0.09 in fourth row of Fig. 6). When *a*_*s*_ increased above grid mesh size, all the fractal designs became excluded by the grid design, irrespective of the environment ruggedness (*a*_*s*_ = 0.33, 0.62 in rows three to five of Fig. 6; Fig. 7, bottom right).

### Pareto-fronts of average rank of errors in problems 1&2

Our results on average rank of errors are reported in Fig. 8. Hybrid designs with small *p*, similar to a grid, reached the lowest average ranks on RRMSE(*ν*) in problem 1 and the lowest average ranks on RRMSE(*γ*) in problem 2. Conversely, fractal designs with intermediate contaction parameter *x* reached the lowest average ranks on RRMSE(*a*_*s*_) in both problems. The global Pareto front on average ranks of errors thus contained both hybrid and fractal designs, emphasizing their complementarity. The steepness of the Pareto front in Fig. 8 shows how much the rank of error on *a*_*s*_ is decreased when increasing the rank of error on *ν* or *γ*. The slope is steeper in problem 2 for a monotonic environment, suggesting that the trade-off is the most interesting in those cases.

**Figure 8.**
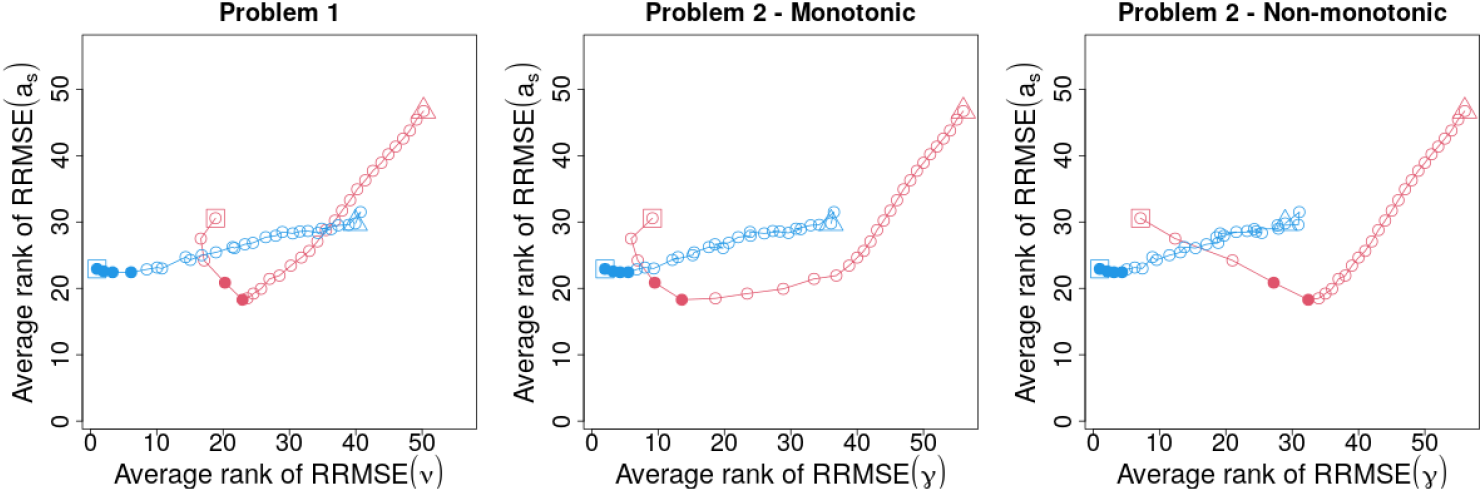
Average rank of designs across the range of *a*_*s*_ values [0.028 − 2.8]. The range corresponds to 0.1 to 10 times the feasible grid mesh size *L/*6. Panel 1 corresponds to problem 1 (mean versus autocorrelation range estimation). Panels 2 and 3 correspond to problem 2 (slope versus autocorrelation range estimation) for monotonic (panel 2) or non monotonic (panel 3) environmental covariate. Red (respectively blue) dots correspond to average rank of RRMSEs for fractal (respectively hybrid) designs with various parameters. Designs with consecutive values of parameters are connected with a line. Triangles indicate most irregular designs within a type. Squares indicate most regular designs within a type. Filled dots show designs that belong to the Pareto front within all designs.

### Minimum distance needed to cover grid and fractal designs

The minimum distance needed to cover fractal designs was between c.a. 50% (lowest *x*) and c.a. 75% (largest *x*) the distance needed to cover the grid design (Fig. 9).

**Figure 9.**
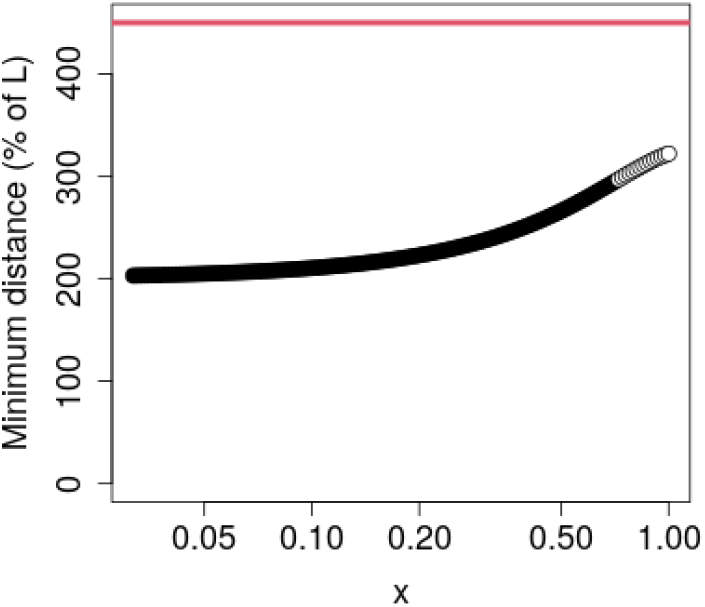
Minimum distance needed to cover fractal and grid design. Black dots show the minimum distance for fractal designs at *x* values where formula (2) is valid. White dots shows the spanning path length of fractal design for *x* values where formula (2) is conservative, and the true shortest path may be even shorter. The horizontal red line shows the spanning path length of the grid design.

## Discussion

Many analyses of species spatial distribution at landscape scale aim at simultaneously estimating effects acting on a populational response variable and the residual spatial autocorrelation range of the response variable (Ranius et al., 2010; Johansson et al., 2012). Here, we compared errors of various sampling designs on both objectives. Our main findings were that: (i) grid design was the best strategy among hybrid designs when averaging over a broad range of possible autocorrelation range values; (ii) when the grid mesh size was markedly larger than the autocorrelation range of the response variable, fractal designs could estimate autocorrelation range more accurately than all the hybrid designs, but their ability to estimate the mean of the response variable or a non-monotonic environmental effect were heavily degraded; (iii) in those situations of small autocorrelation range, fractal designs could outperform hybrid designs at estimating the effect of a monotonic environmental covariate and become the only Pareto-optimal strategies for this type of problem, an advantage that still remained perceivable when averaging over a broader range of autocorrelation ranges; (vi) from a practical perspective, fractal designs may be easier to implement and monitor than a grid design because they have a shorter spanning path.

### Grid design was the best strategy among hybrid designs when averaging over a broad range of possible autocorrelation range values

When autocorrelation range was smaller than the grid mesh size, hybrid designs were spread along a Pareto front between grid and random designs. In other words, increasing the degree of randomness (*p*) in the design tended to degrade the estimation accuracy on the effect but improved the estimation accuracy on the autocorrelation range. This pattern was in line with conclusions of a previous study (Bijleveld et al., 2012), and was robust to changing the target effect (mean or environmental covariate). The fact that grid design was inefficient at quantifying autocorrelation range when mesh size was above the autocorrelation range has been repeatedly emphasized in previous studies, especially in the field of population genetics and isolation by distance studies (Sokal and Oden, 1978; Epperson and Li, 1997). In these situations, pairwise distances smaller than the grid mesh size are needed to accurately estimate the autocorrelation range, and such smaller distances were provided by the introduction of random points in our study, hence leading to the observed Pareto front of hybrid designs.

Conversely, when the autocorrelation range was above the grid mesh size, increasing randomness in hybrid designs always became detrimental to both estimation objectives because accurately estimating effects and autocorrelation range did not require enriching the grid design with small pairwise distances. This finding was robust to the effect considered, leading to the general guideline that grid designs should be favoured when the autocorrelation range of target response variable is larger than the feasible grid mesh size. In line with this advice, a recent simulation study (Perret et al., 2022) focusing on the problem of estimating the mean density of a population (analogous to *ν* in problem 1 here) showed that when grid mesh size is equal to or moderately lower than the autocorrelation range of the density, switching to a random design generated the highest relative increase in error.

Averaging errors across autocorrelation ranges offers a way to assess designs ability to deal with a variety of possible autocorrelation ranges of the response variable. This provides insights about the design suitability for multi-taxonomic empirical studies involving species with contrasted dispersal abilities or home ranges. Using this approach, Bijleveld et al. (2012) found that there was a Pareto front of hybrid designs between grid and random strategies. However, our theoretical analysis showed that the magnitude of errors on autocorrelation range estimation rapidly increases as the autocorrelation range decreases. Direct averaging across aurocorrelation range values thus tends to give higher weights to situations with shorter autocorrelation ranges, where there is indeed a Pareto front of hybrid designs. Here, we controlled this effect by considering error ranks rather than error values when averaging across autocorrelation ranges. The grid design then emerged as the best choice among hybrid designs to deal with a broad range of possible autocorrelation range values. The question that remained was whether fractal designs could show advantages compared to the grid design that hybrid designs did not.

### Fractal designs yielded better estimates of short aurocorrelation ranges than hybrid designs

When autocorrelation range was above the grid mesh size (and not too high to ensure that designs still differed in accuracy), fractal designs were suboptimal compared to grid, because their main contribution was, like random sampling points, to generate small pairwise distances.

When autocorrelation range was lower than grid mesh size, we already mentioned that small pairwise distances between sampling points were useful to estimate the autocorrelation range. When considering autocorrelation ranges for which even the typical pairwise distances generated by random designs are still too large, fractal designs were able to reach lower estimation error of autocorrelation range because decreasing their contraction parameter allowed generating arbitrarily small pairwise distances. This result is typically in line with the study of Ferrandino (2004), which showed that fractals were more efficient at capturing fine scale aggregation of a pathogen distribution than random or grid designs. Therefore, in our study, fractals could constitute new Pareto-optimal strategies, focused on estimating short autocorrelation ranges. Contrary to what could be expected (e.g. Simpson and Pearse (2021)), fractal designs showed no clear pattern of ‘power concentration’ of autocorrelation range estimation (i.e. peaks of accuracy) at few larger scales that would correspond to the design structure. We could visually identify a weak signal of that type for the most irregular fractal designs (Fig. S1), but even in this case it was dampened by the fact that error necessarily increases with scale, as the effective of number pairwise distances available for estimation (i.e. after accounting for pseudo-replication) decreases.

Decreasing the contraction parameter to improve the estimation of short autocorrelation ranges generated a marked agregation of points in space, which degraded a lot the ability to estimate the mean of response variable or the effect of a non-monotonic, rugged environmental covariates. For these problems, choosing fractal designs with low contraction parameters thus amounted to focus exclusively on accurately estimating small autocorrelation ranges, sacrificing other objectives. Under these circumstances, the necessity of keeping a large study area becomes unclear, and reducing the area of study may actually be a better option than implementing a fractal design.

The unbalanced perfomance of fractal designs in estimation problems involving the mean of the response variable or the effect of a non-monotonic covariate became even clearer when considering average errors of designs across a broad range of possible autocorrelation ranges. Some fractal designs could be detected as slightly advantageous, on average, to estimate autocorrelation range, because of their good performance when confronted to short ranges. However, the cost on the estimation error of the effect was quite prohibitive : these designs did not reach the first third of tested options on this criteron.

### Fractal design completely outperformed hybrid designs when studying monotonic covariate under short-ranged autocorrelation

While the interest of fractal designs seemed quite limited when estimating the mean of response variable across the study area or the effect of a rugged environmental covariates, the picture was quite different when studying the effect of a monotonic environmental covariate. There are many empirical questions related to species distributions that involves monotonic environmental gradient in space: the effect of latitudinal gradient and associated climatic trends across a biogeographic region, the effect of altitude on a mountaine slope, the effect of salinity in an estuarine river etc. When studying the effect of this kind of variables, the sparsity of fractal designs turned out to be advantageous, as conjectured by Guo et al. (2023). Moderately increasing the contraction parameter yielded a higher standard deviation of the covariate in the design compared to hybrid designs, hence leading to lower error of the estimated effect. In the meantime, aggregating sampling points in space yielded better estimates of short autocorrelation ranges. Thus, when studying a monotonic covariate under short autocorrelation range, fractal designs with intermediate contraction outperformed hybrid designs on all objectives and constituted the only Pareto-optimal strategies.

This good performance of fractal designs was perceivable in the analysis of average performance of designs across a broader range of autocorrelation range values. Like in other problems, some fractal designs could be detected as slightly advantageous, on average, to estimate autocorrelation range, but this time they also remained in the first third regarding environmental effect estimation, making the trade-off with hybrid designs (and especially the grid) more interesting.

Most interesting fractal designs harboured intermediate contraction parameters, which raises the question of identifying appropriate values in practice. While our aim here is to provide general insights about the relative strength associated to fractal and hybrid designs, precise values should be determined on a case-by-case basis. We strongly encourage operators to adopt a virtual ecology approach (Zurell et al., 2010) and perform *in silico* simulation tests of contraction parameters in their own system with their own budget to find adequate parameter values.

### Sampling budget and sampled area size determine the position of feasible grid mesh size compared to possible autocorrelation ranges

Our conclusions about average performance of designs across possible autocorrelation ranges tightly depend on the range of values over which the average is computed. For instance, if one had only considered autocorrelation range values lower than grid mesh size, a Pareto front of hybrid designs would have emerged rather than the prominence of grid design, while fractal designs would have reached better ranks on their ability to estimate autocorrelation range. Considering autocorrelation range values lower than grid mesh size only would correspond to a sampling budget too low to create a grid with mesh size that matches even the longest autocorrelation ranges. This limitation may arise for large, biogeographic studies of species distributions, but seems quite pessimistic when working at landscape scale over a range of taxa including species with good dispersal abilities or large home ranges. By contrast, considering autocorrelation range values larger than grid mesh size only, would correspond to a sampling budget high enough to create a grid with mesh size smaller than any possible autocorrelation range, which seems quite idealistic. Here, we rather considered a more pragmatical case — likely to correspond to many studies of species distribution within a landscape — where the sampling budget allows implementing a grid design with mesh size taking an intermediate value within possible autocorrelation ranges.

The previous comment shows the critical interplay between the surveyed area and the sampling budget, which co-determine the feasible grid mesh size. This open the question of modulating the area of study, given a sampling budget which we did not explicitly explore here. Reducing the studied area can lead to a shorter grid mesh size. Therefore, if the area is sufficiently reduced, grid design can become lower than most of possible autocorrelation ranges, hence making grid design an unambiguously robust choice. However, the area study cannot be freely reduced because of two limitations. First it increases pseudoreplication for the higher end of possible autocorrelation ranges. Second, it is crucial to keep environmental variation within the area of study when one aims at assessing the effect of environmental gradients (Field et al., 2009; Albert et al., 2010). This is clearly illustrated by our results : the accuracy of the estimate of a covariate effect increases with the standard deviation of the covariate across the area of study (see e.g. our asymptotic analysis of error on *γ* for short autocorrelation range). Reducing the sampling area necessarily contributes to reduce the environmental variation, particularly for monotonic environmental covariates (see Guo et al. (2023) for a discussion on this point).

## Conclusions

Adequate sampling strategies to jointly estimate environmental effects and autocorrelation range of a response variable depend on the number of sampling sites that can be implemented over the study area, and subsequent feasible grid mesh size (Fig. 10). When autocorrelation range values are expected to be above the feasible grid mesh size, the grid design stands out as the best option, and may thus be a robust choice in practical application. By contrast, when autocorrelation range values are expected to be spread around or shorter than the grid mesh size, the choice of designs depends on the effect of interest, and fractal designs outperformed hybrid designs when studying monotonic environmental gradients across the study area. In other situations involving non-monotonic rugged covariates or simply estimating the global mean of target variable over the study area, it seemed more efficient to implement a grid design.

**Figure 10.**
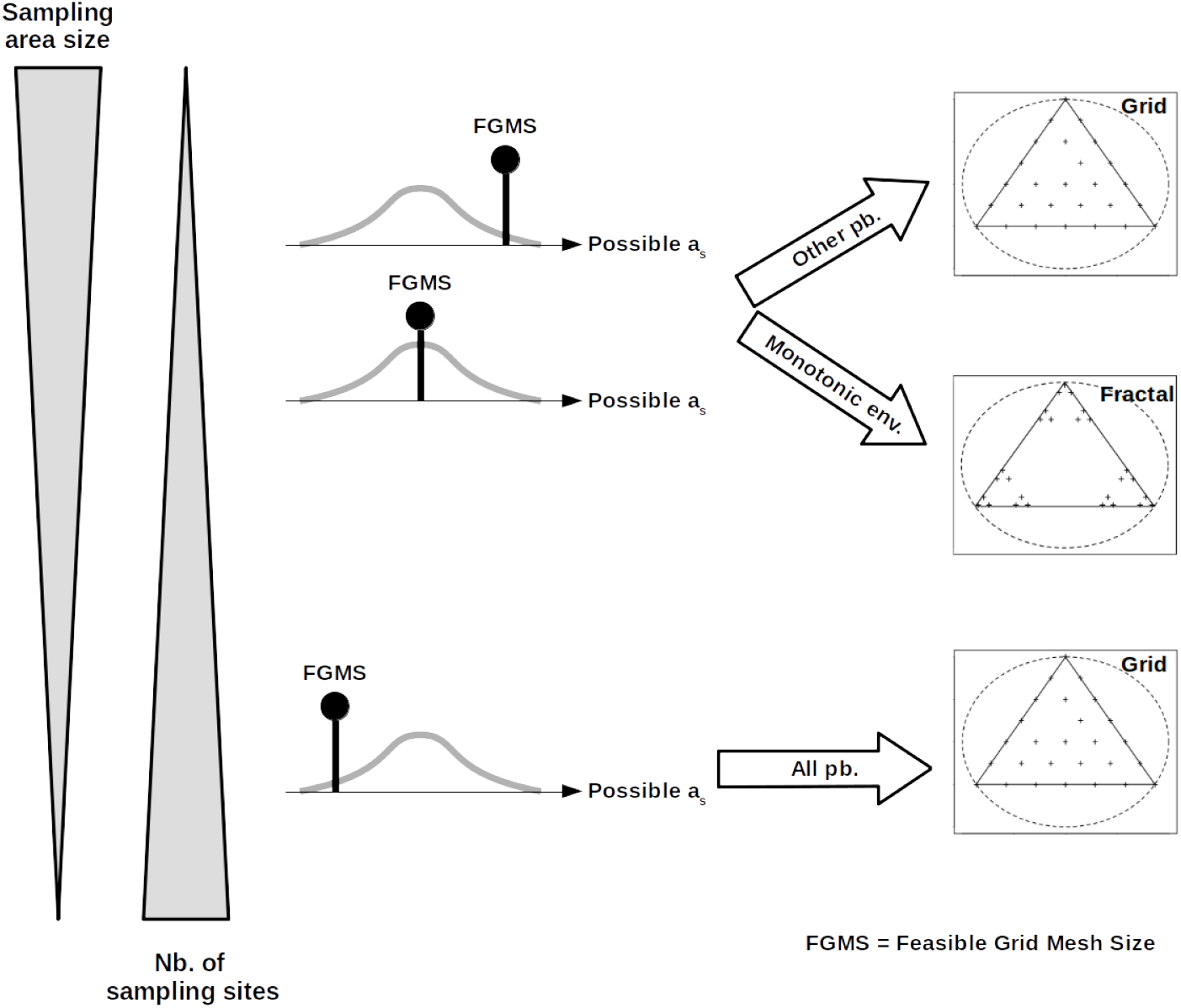
Recommended designs depending on the relative position of the feasible grid mesh size (FGMS) within possible autocorrelation range values. The FGMS decreases with the feasible number of sampling sites (*N* in our study) and increases with sampling area size (e.g. area side length *L* in our study). Conclusions can differ depending on the target estimation ‘problem’. We distinguish cases where, in addition to estimating autocorrelation range, one aims either at estimating the effect of a monotonic environmental covariate across the area (‘Monotonic env.’) or at estimating the effect of a non-monotonic covariate or a constant intercept (‘Other pb.’).

Overall, the niche for fractal designs in terms of estimation error seems limited to estimating the effect of monotonic environmental gradients under autocorrelation ranges that are equal or shorter to feasible grid mesh size. However, it should be noted that estimation errors is one criteria among others to choose a sampling strategy, and it has to be inserted in a broader multi-criteria analysis that also accounts e.g. for the distance needed to perform the survey. On that respect we showed that it is typically shorter to cover a fractal design than a grid design with equivalent sampling budget, which may rehabilitate the fractal option beyond the niche identified here. It should also be noted that we evaluated designs on a simple scenario with a parsimonious autocorrelation structure (only positive autocorrelation, steadily decreasing in space with a single well-defined range). However, biological patterns often stem from heterogeneous drivers acting at different scales (Ricklefs, 2008; Thuiller et al., 2015; Guo et al., 2023). For instance, when studying the distribution of a species in space, competition among conspecifics may drive negative autocorrelation (i.e. underdispersion) at a given spatial scale, while limited dispersal may drive positive autocorrelation at a coarser grain (see Bolker and Pacala (1997) for a theoretical analysis of these effects in plant populations). In the meantime, non-measured environmental variables may also leave various autocorrelation signatures across scales (Legendre, 1993). Designs that harbour a clear hierarchical structure — like fractal designs — may be particularly adapted to capture such heterogeneity (Simpson and Pearse, 2021), provided that the scales of variation induced by the hypothesized processed match the hierarchy of the design.

## Acknowledgements

The author thanks C. Bouget and A. Brin for discussion about the practical implementation of fractal sampling designs, J. Crabot for pointing useful bibliography abour fractal designs, B. Laroche for orienting towards the framework of optimal design, C. Sirami for emphasizing the interest of random designs as a benchmark and C.J. Marsh and N. Yoccoz for their careful reviews of an earlier version of the manuscript.

This research was funded in whole by the Agence Nationale de la Recherche (ANR), through the BloBiForM project grant ANR-19-CE32-0002-01. A CC-BY public copyright license has been applied by the author to the present document and will be applied to all subsequent versions up to the Author Accepted Manuscript arising from this submission, in accordance with the grant’s open access conditions.

## Supplementary figures

**Figure S1:**
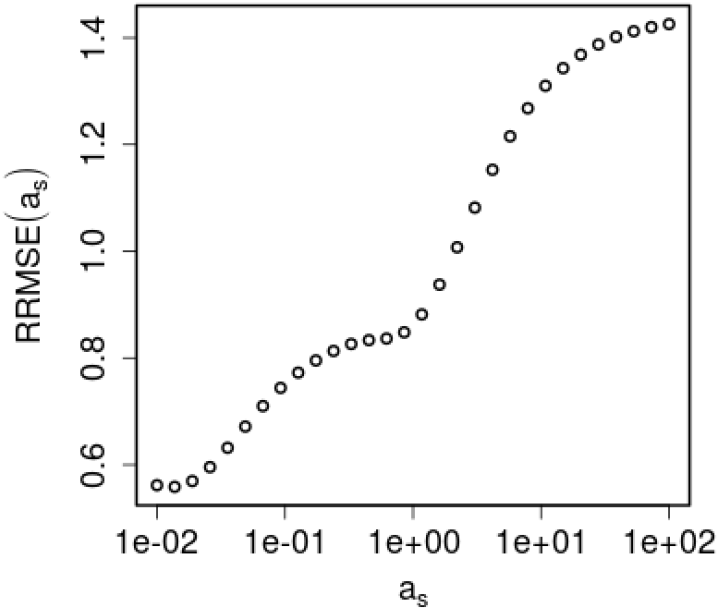
Relative root mean square error of estimation of autocorrelation range (*a*_*s*_) for a fractal design with *x* = 0.035 along a gradient of *a*_*s*_ values. We observe two points of inflexion in the curve : the local minima on the left and a point of zero acceleration in the middle, which both corresponds to effects of power concentration associated to the presence of discrete scale in the design. This type of irregular pattern does not emerge in fractals with larger *x* (i.e. less contrast among scales), nor in hybrid designs (no discrete scales).

