## Appendix S1 for "Efficient sampling designs to assess biodiversity spatial autocorrelation : should we go fractal ?"

### Supplementary information

#### 1 Deriving Fisher information matrix

Let a $N-$sample $\boldsymbol{Y} = (Y_1, ..., Y_N)$ following the statistical model :

$$Y_i = \mu + \beta x_i + Z_i \quad (1)$$

where the vector $\boldsymbol{Z} = (Z_1, ..., Z_N)$ follows a Gaussian Random Field (GRF) with density $f$ on $\mathbb{R}^N$:

$$f(\boldsymbol{z}) = (\det(2\pi\Sigma))^{-\frac{1}{2}} e^{-\frac{1}{2}\boldsymbol{z}'\Sigma^{-1}\boldsymbol{z}}$$

$\Sigma$ is the covariance matrix of the GRF such that:

$$\Sigma_{ij} = \sigma^2 c\left(\frac{d_{ij}}{a_s}\right)$$

with $d_{ij}$ the euclidean distance between points $i$ and $j$, $\sigma^2$ the variance of observation at each point, $c(.)$ a correlation function which decreases from 1 to 0 over $\mathbb{R}^+$ and $a_s$ a scaling factor. We note $\tilde{\Sigma}$ the correlation matrix associated to covariance matrix $\Sigma$

The log-likelihood of parameters is:

$$l(\mu, \beta, \sigma^2, a_s|\boldsymbol{z}) = -\frac{1}{2}\left(N\log(2\pi) + \log(\det\Sigma) + (\boldsymbol{y} - \mu\mathbf{1} - \beta\boldsymbol{x})'\Sigma^{-1}(\boldsymbol{y} - \mu\mathbf{1} - \beta\boldsymbol{x})\right)$$

Then:

$$\begin{aligned}
\frac{\partial l}{\partial \mu} &= \mathbf{1}'\Sigma^{-1}(\boldsymbol{y} - \mu\mathbf{1} - \beta\boldsymbol{x}) \\
\frac{\partial l}{\partial \beta} &= \boldsymbol{x}'\Sigma^{-1}(\boldsymbol{y} - \mu\mathbf{1} - \beta\boldsymbol{x}) \\
\frac{\partial l}{\partial \sigma^2} &= -\frac{1}{2}\left(\frac{1}{\det\Sigma}\frac{\partial \det\Sigma}{\partial \sigma^2} + (\boldsymbol{y} - \mu\mathbf{1} - \beta\boldsymbol{x})'\frac{\partial \Sigma^{-1}}{\partial \sigma^2}(\boldsymbol{y} - \mu\mathbf{1} - \beta\boldsymbol{x})\right) \\
\frac{\partial l}{\partial a_s} &= -\frac{1}{2}\left(\frac{1}{\det\Sigma}\frac{\partial \det\Sigma}{\partial a_s} + (\boldsymbol{y} - \mu\mathbf{1} - \beta\boldsymbol{x})'\frac{\partial \Sigma^{-1}}{\partial a_s}(\boldsymbol{y} - \mu\mathbf{1} - \beta\boldsymbol{x})\right)
\end{aligned}$$

Recalling Jacobi's formula :

$$\frac{\partial \det A}{\partial \theta} = (\det A)\,\mathrm{tr}\left(A^{-1}\frac{\partial A}{\partial \theta}\right)$$

one obtains that:

$$\begin{aligned}
\frac{\partial l}{\partial \mu} &= \mathbf{1}'\Sigma^{-1}(\boldsymbol{y} - \mu\mathbf{1} - \beta\boldsymbol{x}) \\
\frac{\partial l}{\partial \beta} &= \boldsymbol{x}'\Sigma^{-1}(\boldsymbol{y} - \mu\mathbf{1} - \beta\boldsymbol{x}) \\
\frac{\partial l}{\partial \sigma^2} &= -\frac{1}{2}\left(\mathrm{tr}(\Sigma^{-1}\frac{\partial \Sigma}{\partial \sigma^2}) + (\boldsymbol{y} - \mu\mathbf{1} - \beta\boldsymbol{x})'\frac{\partial \Sigma^{-1}}{\partial \sigma^2}(\boldsymbol{y} - \mu\mathbf{1} - \beta\boldsymbol{x})\right) \\
\frac{\partial l}{\partial a_s} &= -\frac{1}{2}\left(\mathrm{tr}(\Sigma^{-1}\frac{\partial \Sigma}{\partial a_s}) + (\boldsymbol{y} - \mu\mathbf{1} - \beta\boldsymbol{x})'\frac{\partial \Sigma^{-1}}{\partial a_s}(\boldsymbol{y} - \mu\mathbf{1} - \beta\boldsymbol{x})\right)
\end{aligned}$$

Then one can derive second order derivatives:

$$
\begin{aligned}
\frac{\partial^2 l}{(\partial\mu)^2} &= -\mathbf{1}'\Sigma^{-1}\mathbf{1} \\
\frac{\partial^2 l}{\partial\mu\partial\beta} &= -\mathbf{1}'\Sigma^{-1}\boldsymbol{x} \\
\frac{\partial^2 l}{\partial\mu\partial\sigma^2} &= \mathbf{1}'\frac{\partial\Sigma^{-1}}{\partial\sigma^2}\left(\boldsymbol{y}-\mu\mathbf{1}-\beta\boldsymbol{x}\right) \\
\frac{\partial^2 l}{\partial\mu\partial a_s} &= \mathbf{1}'\frac{\partial\Sigma^{-1}}{\partial a_s}\left(\boldsymbol{y}-\mu\mathbf{1}-\beta\boldsymbol{x}\right) \\
\frac{\partial^2 l}{(\partial\beta)^2} &= -\boldsymbol{x}'\Sigma^{-1}\boldsymbol{x} \\
\frac{\partial^2 l}{\partial\beta\partial\sigma^2} &= \boldsymbol{x}'\frac{\partial\Sigma^{-1}}{\partial\sigma^2}\left(\boldsymbol{y}-\mu\mathbf{1}-\beta\boldsymbol{x}\right) \\
\frac{\partial^2 l}{\partial\beta\partial a_s} &= \boldsymbol{x}'\frac{\partial\Sigma^{-1}}{\partial a_s}\left(\boldsymbol{y}-\mu\mathbf{1}-\beta\boldsymbol{x}\right) \\
\frac{\partial^2 l}{(\partial\sigma^2)^2} &= -\frac{1}{2}\left(\mathrm{tr}\left(\frac{\partial\Sigma^{-1}}{\partial\sigma^2}\frac{\partial\Sigma}{\partial\sigma^2}+\Sigma^{-1}\frac{\partial^2\Sigma}{(\partial\sigma^2)^2}\right)+\left(\boldsymbol{y}-\mu\mathbf{1}-\beta\boldsymbol{x}\right)'\frac{\partial^2\Sigma^{-1}}{(\partial\sigma^2)^2}\left(\boldsymbol{y}-\mu\mathbf{1}-\beta\boldsymbol{x}\right)\right) \\
\frac{\partial^2 l}{\partial\sigma^2\partial a_s} &= -\frac{1}{2}\left(\mathrm{tr}\left(\frac{\partial\Sigma^{-1}}{\partial a_s}\frac{\partial\Sigma}{\partial\sigma^2}+\Sigma^{-1}\frac{\partial^2\Sigma}{\partial\sigma^2\partial a_s}\right)+\left(\boldsymbol{y}-\mu\mathbf{1}-\beta\boldsymbol{x}\right)'\frac{\partial^2\Sigma^{-1}}{\partial\sigma^2\partial a_s}\left(\boldsymbol{y}-\mu\mathbf{1}-\beta\boldsymbol{x}\right)\right) \\
\frac{\partial^2 l}{(\partial a_s)^2} &= -\frac{1}{2}\left(\mathrm{tr}\left(\frac{\partial\Sigma^{-1}}{\partial a_s}\frac{\partial\Sigma}{\partial a_s}+\Sigma^{-1}\frac{\partial^2\Sigma}{(\partial a_s)^2}\right)+\left(\boldsymbol{y}-\mu\mathbf{1}-\beta\boldsymbol{x}\right)'\frac{\partial^2\Sigma^{-1}}{(\partial a_s)^2}\left(\boldsymbol{y}-\mu\mathbf{1}-\beta\boldsymbol{x}\right)\right)
\end{aligned}
$$

Taking probabilistic expectation of these second derivatives over the random variable $\boldsymbol{Y}$ at true parameters and using that:

$$\mathbb{E}\left[(\boldsymbol{Y}-\mu\mathbf{1}-\beta\boldsymbol{x})'A(\boldsymbol{Y}-\mu\mathbf{1}-\beta\boldsymbol{x})\right]=\mathrm{tr}(\Sigma A)$$

one obtains that:

$$
\begin{aligned}
\mathbb{E}\left[\frac{\partial^2 l}{(\partial\mu)^2}\right] &= -\mathbf{1}'\Sigma^{-1}\mathbf{1} \\
\mathbb{E}\left[\frac{\partial^2 l}{\partial\mu\partial\beta}\right] &= -\mathbf{1}'\Sigma^{-1}\boldsymbol{x} \\
\mathbb{E}\left[\frac{\partial^2 l}{\partial\mu\partial\sigma^2}\right] &= 0 \\
\mathbb{E}\left[\frac{\partial^2 l}{\partial\mu\partial a_s}\right] &= 0 \\
\mathbb{E}\left[\frac{\partial^2 l}{(\partial\beta)^2}\right] &= -\boldsymbol{x}'\Sigma^{-1}\boldsymbol{x} \\
\mathbb{E}\left[\frac{\partial^2 l}{\partial\beta\partial\sigma^2}\right] &= 0 \\
\mathbb{E}\left[\frac{\partial^2 l}{\partial\beta\partial a_s}\right] &= 0 \\
\mathbb{E}\left[\frac{\partial^2 l}{(\partial\sigma^2)^2}\right] &= -\frac{1}{2}\left(\mathrm{tr}\left(\frac{\partial\Sigma^{-1}}{\partial\sigma^2}\frac{\partial\Sigma}{\partial\sigma^2}+\Sigma^{-1}\frac{\partial^2\Sigma}{(\partial\sigma^2)^2}+\Sigma\frac{\partial^2\Sigma^{-1}}{(\partial\sigma^2)^2}\right)\right) \\
\mathbb{E}\left[\frac{\partial^2 l}{\partial\sigma^2\partial a_s}\right] &= -\frac{1}{2}\left(\mathrm{tr}\left(\frac{\partial\Sigma^{-1}}{\partial a_s}\frac{\partial\Sigma}{\partial\sigma^2}+\Sigma^{-1}\frac{\partial^2\Sigma}{\partial\sigma^2\partial a_s}+\Sigma\frac{\partial^2\Sigma^{-1}}{\partial\sigma^2\partial a_s}\right)\right) \\
\mathbb{E}\left[\frac{\partial^2 l}{(\partial a_s)^2}\right] &= -\frac{1}{2}\left(\mathrm{tr}\left(\frac{\partial\Sigma^{-1}}{\partial a_s}\frac{\partial\Sigma}{\partial a_s}+\Sigma^{-1}\frac{\partial^2\Sigma}{(\partial a_s)^2}+\Sigma\frac{\partial^2\Sigma^{-1}}{(\partial a_s)^2}\right)\right)
\end{aligned}
$$

Noting that:

$$0=\frac{\partial\Sigma^{-1}\Sigma}{\partial\theta_1}=\frac{\partial\Sigma^{-1}}{\partial\theta_1}\Sigma+\Sigma^{-1}\frac{\partial\Sigma}{\partial\theta_1}$$

and that :

$$0=\frac{\partial^2\Sigma^{-1}\Sigma}{\partial\theta_1\partial\theta_2}=\frac{\partial^2\Sigma^{-1}}{\partial\theta_1\partial\theta_2}\Sigma+\frac{\partial\Sigma^{-1}}{\partial\theta_1}\frac{\partial\Sigma}{\partial\theta_2}+\frac{\partial\Sigma^{-1}}{\partial\theta_2}\frac{\partial\Sigma}{\partial\theta_1}+\Sigma^{-1}\frac{\partial^2\Sigma}{\partial\theta_1\partial\theta_2}$$

the three last second order derivatives can be simplified into:

$$\begin{array}{rcl}
\mathbb{E}\left[\frac{\partial^2 l}{(\partial\sigma^2)^2}\right] & = & \frac{1}{2}\mathrm{tr}\left(\frac{\partial\Sigma^{-1}}{\partial\sigma^2}\frac{\partial\Sigma}{\partial\sigma^2}\right) = -\frac{1}{2}\mathrm{tr}\left(\Sigma^{-1}\frac{\partial\Sigma}{\partial\sigma^2}\Sigma^{-1}\frac{\partial\Sigma}{\partial\sigma^2}\right) \\
\mathbb{E}\left[\frac{\partial^2 l}{\partial\sigma^2\partial a_s}\right] & = & \frac{1}{2}\mathrm{tr}\left(\frac{\partial\Sigma^{-1}}{\partial\sigma^2}\frac{\partial\Sigma}{\partial a_s}\right) = -\frac{1}{2}\mathrm{tr}\left(\Sigma^{-1}\frac{\partial\Sigma}{\partial\sigma^2}\Sigma^{-1}\frac{\partial\Sigma}{\partial a_s}\right) \\
\mathbb{E}\left[\frac{\partial^2 l}{(\partial a_s)^2}\right] & = & \frac{1}{2}\mathrm{tr}\left(\frac{\partial\Sigma^{-1}}{\partial a_s}\frac{\partial\Sigma}{\partial a_s}\right) = -\frac{1}{2}\mathrm{tr}\left(\Sigma^{-1}\frac{\partial\Sigma}{\partial a_s}\Sigma^{-1}\frac{\partial\Sigma}{\partial a_s}\right)
\end{array}$$

Further using that:

$$\frac{\partial\Sigma}{\partial\sigma^2} = \frac{1}{\sigma^2}\Sigma$$

one obtains that :

$$\begin{array}{rcl}
\mathbb{E}\left[\frac{\partial^2 l}{(\partial\sigma^2)^2}\right] & = & -\frac{N}{2\sigma^4} \\
\mathbb{E}\left[\frac{\partial^2 l}{\partial\sigma^2\partial a_s}\right] & = & -\frac{1}{2\sigma^2}\mathrm{tr}\left(\Sigma^{-1}\frac{\partial\Sigma}{\partial a_s}\right) = -\frac{1}{2\sigma^2}\mathrm{tr}\left(\tilde{\Sigma}^{-1}\frac{\partial\tilde{\Sigma}}{\partial a_s}\right) \\
\mathbb{E}\left[\frac{\partial^2 l}{(\partial a_s)^2}\right] & = & -\frac{1}{2}\mathrm{tr}\left(\Sigma^{-1}\frac{\partial\Sigma}{\partial a_s}\Sigma^{-1}\frac{\partial\Sigma}{\partial a_s}\right) = -\frac{1}{2}\mathrm{tr}\left(\tilde{\Sigma}^{-1}\frac{\partial\tilde{\Sigma}}{\partial a_s}\tilde{\Sigma}^{-1}\frac{\partial\tilde{\Sigma}}{\partial a_s}\right)
\end{array}$$

Therefore, the Fisher information matrix of model (1) is:

$$\mathcal{I}(\mu,\beta,\sigma^2,a_s) = \\ \begin{pmatrix}
\mathbf{1}'\Sigma^{-1}\mathbf{1} & \mathbf{1}'\Sigma^{-1}\boldsymbol{x} & 0 & 0 \\
\mathbf{1}'\Sigma^{-1}\boldsymbol{x} & \boldsymbol{x}'\Sigma^{-1}\boldsymbol{x} & 0 & 0 \\
0 & 0 & \frac{N}{2\sigma^4} & \frac{1}{2\sigma^2}\mathrm{tr}(\tilde{\Sigma}^{-1}\frac{\partial\tilde{\Sigma}}{\partial a_s}) \\
0 & 0 & \frac{1}{2\sigma^2}\mathrm{tr}(\tilde{\Sigma}^{-1}\frac{\partial\tilde{\Sigma}}{\partial a_s}) & \frac{1}{2}\mathrm{tr}(\tilde{\Sigma}^{-1}\frac{\partial\tilde{\Sigma}}{\partial a_s}\tilde{\Sigma}^{-1}\frac{\partial\tilde{\Sigma}}{\partial a_s})
\end{pmatrix} \quad (2)$$

**Problem 1: mean versus autocorrelation range estimation**

Recall the statistical model associated to problem 1:

$$Y_i = \mu + Z_i$$

which is a simplification of model (1), obtained by taking $\beta = 0$. The associated Fisher matrix can be directly derived from (2) as:

$$\mathcal{I}(\mu,\sigma^2,a_s) = \begin{pmatrix}
\mathbf{1}'\Sigma^{-1}\mathbf{1} & 0 & 0 \\
0 & \frac{N}{2\sigma^4} & \frac{1}{2\sigma^2}\mathrm{tr}(\tilde{\Sigma}^{-1}\frac{\partial\tilde{\tilde{\Sigma}}}{\partial a_s}) \\
0 & \frac{1}{2\sigma^2}\mathrm{tr}(\tilde{\Sigma}^{-1}\frac{\partial\tilde{\Sigma}}{\partial a_s}) & \frac{1}{2}\mathrm{tr}(\tilde{\Sigma}^{-1}\frac{\partial\tilde{\Sigma}}{\partial a_s}\tilde{\Sigma}^{-1}\frac{\partial\tilde{\Sigma}}{\partial a_s})
\end{pmatrix}$$

Changing parameters $\mu$ to exponential parameters $\nu = e^{\mu}$ and defining the vector of parameter $\boldsymbol{\theta} = (\nu, \sigma^2, a_s)$ yields the Fisher information matrix :

$$\mathcal{I}(\boldsymbol{\theta}) = \begin{pmatrix}
\frac{1}{\nu^2}\mathbf{1}'\Sigma^{-1}\mathbf{1} & 0 & 0 \\
0 & \frac{N}{2\sigma^4} & \frac{1}{2\sigma^2}\mathrm{tr}(\tilde{\Sigma}^{-1}\frac{\partial\tilde{\tilde{\Sigma}}}{\partial a_s}) \\
0 & \frac{1}{2\sigma^2}\mathrm{tr}(\tilde{\Sigma}^{-1}\frac{\partial\tilde{\Sigma}}{\partial a_s}) & \frac{1}{2}\mathrm{tr}(\tilde{\Sigma}^{-1}\frac{\partial\tilde{\Sigma}}{\partial a_s}\tilde{\Sigma}^{-1}\frac{\partial\tilde{\Sigma}}{\partial a_s})
\end{pmatrix} \quad (3)$$

From the Fisher information matrix, one can derive the expected mean squared errors associated to $\hat{\nu}$ and $\hat{a}_s$, denoted MSE($\nu$) and MSE($a_s$) respectively:

$$\mathrm{MSE}(\nu) = \nu^2\sigma^2 \frac{1}{\mathbf{1}'\tilde{\Sigma}^{-1}\mathbf{1}}$$

$$\mathrm{MSE}(a_s) = \frac{2}{\mathrm{tr}(\tilde{\Sigma}^{-1}\frac{\partial\tilde{\Sigma}}{\partial a_s}\tilde{\Sigma}^{-1}\frac{\partial\tilde{\Sigma}}{\partial a_s}) - \frac{1}{N}\mathrm{tr}(\tilde{\Sigma}^{-1}\frac{\partial\tilde{\Sigma}}{\partial a_s})\mathrm{tr}(\tilde{\Sigma}^{-1}\frac{\partial\tilde{\Sigma}}{\partial a_s})}$$

where we used $\Sigma = \sigma^2\tilde{\Sigma}$. Using above expression, the derivation of corresponding RRMSEs is straightforward.

**Problem 2: slope of environmental effect versus autocorrelation range estimation**

Recall the statistical model associated to problem 2:

$$Y_i = \tilde{\mu} + \beta\tilde{x}_i + Z_i$$

which is identical to model (1) up to notational changes. The Fisher matrix of this model is thus directly obtained from (2):

$$\mathcal{I}(\tilde{\mu}, \beta, \sigma^2, a_s) = \begin{pmatrix} \mathbf{1}'\Sigma^{-1}\mathbf{1} & \mathbf{1}'\Sigma^{-1}\tilde{\boldsymbol{x}} & 0 & 0 \\ \mathbf{1}'\Sigma^{-1}\tilde{\boldsymbol{x}} & \tilde{\boldsymbol{x}}'\Sigma^{-1}\tilde{\boldsymbol{x}} & 0 & 0 \\ 0 & 0 & \frac{N}{2\sigma^4} & \frac{1}{2\sigma^2}\mathrm{tr}(\tilde{\Sigma}^{-1}\frac{\partial\tilde{\Sigma}}{\partial a_s}) \\ 0 & 0 & \frac{1}{2\sigma^2}\mathrm{tr}(\tilde{\Sigma}^{-1}\frac{\partial\tilde{\Sigma}}{\partial a_s}) & \frac{1}{2}\mathrm{tr}(\tilde{\Sigma}^{-1}\frac{\partial\tilde{\Sigma}}{\partial a_s}\tilde{\Sigma}^{-1}\frac{\partial\tilde{\Sigma}}{\partial a_s}) \end{pmatrix}$$

Changing parameters $\beta$ to exponential parameters $\gamma = e^\beta$ and defining the vector of parameter $\boldsymbol{\theta} = (\tilde{\mu}, \gamma, \sigma^2, a_s)$ yields the Fisher information matrix :

$$\mathcal{I}(\boldsymbol{\theta}) = \begin{pmatrix} \mathbf{1}'\Sigma^{-1}\mathbf{1} & \frac{1}{\gamma}\mathbf{1}'\Sigma^{-1}\tilde{\boldsymbol{x}} & 0 & 0 \\ \frac{1}{\gamma}\mathbf{1}'\Sigma^{-1}\tilde{\boldsymbol{x}} & \frac{1}{\gamma^2}\tilde{\boldsymbol{x}}'\Sigma^{-1}\tilde{\boldsymbol{x}} & 0 & 0 \\ 0 & 0 & \frac{N}{2\sigma^4} & \frac{1}{2\sigma^2}\mathrm{tr}(\tilde{\Sigma}^{-1}\frac{\partial\tilde{\tilde{\Sigma}}}{\partial a_s}) \\ 0 & 0 & \frac{1}{2\sigma^2}\mathrm{tr}(\tilde{\Sigma}^{-1}\frac{\partial\tilde{\Sigma}}{\partial a_s}) & \frac{1}{2}\mathrm{tr}(\tilde{\Sigma}^{-1}\frac{\partial\tilde{\Sigma}}{\partial a_s}\tilde{\Sigma}^{-1}\frac{\partial\tilde{\Sigma}}{\partial a_s}) \end{pmatrix} \tag{4}$$

From the Fisher information matrix, one can derive the expected mean squared errors associated to $\hat{\gamma}$ and $\hat{a}_s$, denoted MSE($\gamma$) and MSE($a_s$) respectively:

$$\mathrm{MSE}(\gamma) = \gamma^2\sigma^2 \frac{\mathbf{1}'\tilde{\Sigma}^{-1}\mathbf{1}}{(\tilde{\boldsymbol{x}}'\tilde{\Sigma}^{-1}\tilde{\boldsymbol{x}})(\mathbf{1}'\tilde{\Sigma}^{-1}\mathbf{1}) - (\mathbf{1}'\tilde{\Sigma}^{-1}\tilde{\boldsymbol{x}})^2}$$

$$\mathrm{MSE}(a_s) = \frac{2}{\mathrm{tr}(\tilde{\Sigma}^{-1}\frac{\partial\tilde{\Sigma}}{\partial a_s}\tilde{\Sigma}^{-1}\frac{\partial\tilde{\Sigma}}{\partial a_s}) - \frac{1}{N}\mathrm{tr}(\tilde{\Sigma}^{-1}\frac{\partial\tilde{\Sigma}}{\partial a_s})\mathrm{tr}(\tilde{\Sigma}^{-1}\frac{\partial\tilde{\Sigma}}{\partial a_s})}$$

Using above expression, the derivation of corresponding RRMSEs is straightforward.

### 2 Asymptotic errors when $a_s \to 0, +\infty$

#### Exponential mean in problem 1

Recall that $\mathrm{RRMSE}(\nu) = \sigma/\sqrt{\mathbf{1}'\tilde{\Sigma}^{-1}\mathbf{1}}$.

**In the limit $a_s \to 0$:** $\tilde{\Sigma} \to I$, where $I$ stand for the identity matrix. Therefore $\tilde{\Sigma}^{-1} \to I$ and $\mathbf{1}'\tilde{\Sigma}^{-1}\mathbf{1} \to N$, leading in turn to:

$$\text{RRMSE}(\nu) \to \sigma/\sqrt{N}$$

**In the limit $a_s \to +\infty$:** we note $\tilde{\Sigma} = J - \delta E$ where $J_{ij} = 1$ for all $i, j$, $E$ is an invertible matrix and $\delta \to 0^+$. The inverse of $\tilde{\Sigma}$ is:

$$\tilde{\Sigma}^{-1} = -\delta^{-1}E^{-1}\left(I + \frac{\delta^{-1}JE^{-1}}{1 - \delta^{-1}\mathbf{1}'E^{-1}\mathbf{1}}\right)$$

and:

$$\mathbf{1}'\tilde{\Sigma}^{-1}\mathbf{1} = -\delta^{-1}\left(\mathbf{1}'E^{-1}\mathbf{1} + \frac{\delta^{-1}\mathbf{1}'E^{-1}JE^{-1}}{1 - \delta^{-1}\mathbf{1}'E^{-1}\mathbf{1}}\mathbf{1}\right) = \frac{\delta^{-1}\mathbf{1}'E^{-1}\mathbf{1}}{\delta^{-1}\mathbf{1}'E^{-1}\mathbf{1} - 1} \to 1$$

Therefore:

$$\text{RRMSE}(\nu) \to \sigma$$

#### Exponential slope in problem 2

Recall that $\text{RRMSE}(\gamma) = \sigma\sqrt{\mathbf{1}'\tilde{\Sigma}^{-1}\mathbf{1}}/\sqrt{(\tilde{\boldsymbol{x}}'\tilde{\Sigma}^{-1}\tilde{\boldsymbol{x}})(\mathbf{1}'\tilde{\Sigma}^{-1}\mathbf{1}) - (\mathbf{1}'\tilde{\Sigma}^{-1}\tilde{\boldsymbol{x}})^2}$.

**In the limit $a_s \to 0$:** $\mathbf{1}'\tilde{\Sigma}^{-1}\mathbf{1} \to N$, $\mathbf{1}'\tilde{\Sigma}^{-1}\tilde{\boldsymbol{x}} \to 0$ and $\boldsymbol{x}'\tilde{\Sigma}^{-1}\tilde{\boldsymbol{x}} \to N\text{Var}(\tilde{\boldsymbol{x}})$, leading in turn to:

$$\text{RRMSE}(\gamma) \to \frac{\sigma}{SD(\tilde{\boldsymbol{x}})\sqrt{N}}$$

**In the limit $a_s \to +\infty$:** we use, like for exponential mean in problem 1, $\tilde{\Sigma} = J - \delta E$. Then :

$$\mathbf{1}'\tilde{\Sigma}^{-1}\boldsymbol{x} = -\delta^{-1}\mathbf{1}'E^{-1}\boldsymbol{x} + \frac{-\delta^{-2}\mathbf{1}'E^{-1}\mathbf{1}\mathbf{1}'E^{-1}\boldsymbol{x}}{1 - \delta^{-1}\mathbf{1}'E^{-1}\mathbf{1}} = \frac{\delta^{-1}\mathbf{1}'E^{-1}\boldsymbol{x}}{\delta^{-1}\mathbf{1}'E^{-1}\mathbf{1} - 1} \to \frac{\mathbf{1}'E^{-1}\boldsymbol{x}}{\mathbf{1}'E^{-1}\mathbf{1}}$$

and:

$$\boldsymbol{x}'\tilde{\Sigma}^{-1}\boldsymbol{x} = -\delta^{-1}\boldsymbol{x}'E^{-1}\boldsymbol{x} + \frac{-\delta^{-2}\boldsymbol{x}'E^{-1}\mathbf{1}\mathbf{1}'E^{-1}\boldsymbol{x}}{1 - \delta^{-1}\mathbf{1}'E^{-1}\mathbf{1}} \sim \delta^{-1}\left(\frac{(\mathbf{1}'E^{-1}\boldsymbol{x})^2}{\mathbf{1}'E^{-1}\mathbf{1}} - \boldsymbol{x}'E^{-1}\boldsymbol{x}\right)$$

Therefore:

$$\text{RRMSE}(\gamma) \to 0$$

#### Autocorrelation range

Here we consider the particular case of the exponential covariance model : $c(x) = e^{-x}$. Then $\frac{\partial \Sigma}{\partial a_s} = \frac{\sigma^2}{a_s^2} D \circ \Sigma$ where $D$ is the distance matrix among sampling points and $\circ$ is the Hadamard product.

**When** $a_s \to 0$

$$\frac{a_s^2}{d_{\min}} e^{\frac{d_{\min}}{a_s}} \frac{\partial \Sigma}{\partial a_s} \to \sigma^2 R$$
$$e^{\frac{d_{\min}}{a_s}} (I - \sigma^2 \Sigma^{-1}) \to R$$

where $d_{\min}$ is the smallest distance between two distinct sampling points and $R_{ij} = 1$ if $D_{ij} = d_{\min}$, 0 otherwise. Then, $\frac{a_s^2}{d_{min}} e^{\frac{2d_{min}}{a_s}} (\frac{\partial \Sigma}{\partial a_s} - \sigma^2 \Sigma^{-1} \frac{\partial \Sigma}{\partial a_s}) \to \sigma^2 R^2$ which implies in turn

$$\frac{a_s^2}{d_{min}} e^{\frac{2d_{min}}{a_s}} \mathrm{tr}\left(\Sigma^{-1} \frac{\partial \Sigma}{\partial a_s}\right) \to -\mathrm{tr}(R^2)$$

In addition,

$$\begin{aligned}
\mathrm{tr}\left(\Sigma^{-1}\frac{\partial \Sigma}{\partial a_s}\Sigma^{-1}\frac{\partial \Sigma}{\partial a_s}\right) &= \frac{1}{\sigma^4}\mathrm{tr}\left(\sigma^2\Sigma^{-1}\frac{\partial \Sigma}{\partial a_s}\sigma^2\Sigma^{-1}\frac{\partial \Sigma}{\partial a_s}\right) \\
&= \frac{1}{\sigma^4}\mathrm{tr}\left((\sigma^2\Sigma^{-1} - I)\frac{\partial \Sigma}{\partial a_s}\sigma^2\Sigma^{-1}\frac{\partial \Sigma}{\partial a_s} + \frac{\partial \Sigma}{\partial a_s}\sigma^2\Sigma^{-1}\frac{\partial \Sigma}{\partial a_s}\right) \\
&= \frac{1}{\sigma^4}\mathrm{tr}\begin{pmatrix} (\sigma^2\Sigma^{-1} - I)\frac{\partial \Sigma}{\partial a_s}(\sigma^2\Sigma^{-1} - I)\frac{\partial \Sigma}{\partial a_s} \\ +\frac{\partial \Sigma}{\partial a_s}\sigma^2\Sigma^{-1}\frac{\partial \Sigma}{\partial a_s} \\ +(\sigma^2\Sigma^{-1} - I)\left(\frac{\partial \Sigma}{\partial a_s}\right)^2 \end{pmatrix} \\
&= \frac{1}{\sigma^4}\mathrm{tr}\begin{pmatrix} (\sigma^2\Sigma^{-1} - I)\frac{\partial \Sigma}{\partial a_s}(\sigma^2\Sigma^{-1} - I)\frac{\partial \Sigma}{\partial a_s} \\ +\frac{\partial \Sigma}{\partial a_s}(\sigma^2\Sigma^{-1} - I)\frac{\partial \Sigma}{\partial a_s} \\ +(\sigma^2\Sigma^{-1} - I)\left(\frac{\partial \Sigma}{\partial a_s}\right)^2 \\ +\left(\frac{\partial \Sigma}{\partial a_s}\right)^2 \end{pmatrix}
\end{aligned}$$

Therefore,

$$\frac{a_s^4}{d_{\min}^2} e^{\frac{2d_{\min}}{a_s}} \mathrm{tr}\left(\Sigma^{-1}\frac{\partial \Sigma}{\partial a_s}\Sigma^{-1}\frac{\partial \Sigma}{\partial a_s}\right) \to \mathrm{tr}(R^2)$$

which implies that :

$$\lim_{a_s \to 0}\left(\frac{d_{\min}^2}{a_s^4} e^{-\frac{2d_{\min}}{a_s}} \mathrm{MSE}(a_s)\right) = \frac{2}{\mathrm{tr}(R^2)}$$

In other words, $\mathrm{MSE}(a_s)$ increases towards $+\infty$ like $a_s^4 e^{\frac{2d_{\min}}{a_s}}$ when $a_s$ decreases towards 0.

**When** $a_s \to +\infty$ The assumption of exponential covariance implies that $a_s E \to D$ (see previous section for $E$ definition), from which one obtains that $\frac{1}{a_s}E^{-1} \to D^{-1}$. Recalling that $\Sigma^{-1} = \frac{1}{\sigma^2 \mathbb{S}(E^{-1}) - 1}E^{-1}JE^{-1} - \frac{1}{\sigma^2}E^{-1}$, one obtains that:

$$a_s^2 \frac{\partial \Sigma}{\partial a_s} \to \sigma^2 D \circ J = \sigma^2 D$$
$$\frac{1}{a_s}\Sigma^{-1} \to \frac{1}{\sigma^2}\left(\frac{1}{\mathbb{S}(D^{-1})}D^{-1}JD^{-1} - D^{-1}\right)$$

which implies:

$$\begin{array}{lll} a_s \mathrm{tr}\left(\Sigma^{-1}\frac{\partial \Sigma}{\partial a_s}\right) & \rightarrow & \frac{1}{\mathbb{S}(D^{-1})}\mathrm{tr}\left(D^{-1}J\right) - N = 1 - N \\ a_s^2 \mathrm{tr}\left(\Sigma^{-1}\frac{\partial \Sigma}{\partial a_s}\Sigma^{-1}\frac{\partial \Sigma}{\partial a_s}\right) & \rightarrow & \mathrm{tr}\left(\left(\frac{1}{\mathbb{S}(D^{-1})}D^{-1}J - I\right)\left(\frac{1}{\mathbb{S}(D^{-1})}D^{-1}J - I\right)\right) = N - 1 \end{array}$$

Therefore :

$$\lim_{a_s \to +\infty} \frac{1}{a_s^2}\mathrm{MSE}(a_s) = \frac{2N}{N-1}$$

In other words, $\mathrm{MSE}(a_s)$ increases towards $+\infty$ like $a_s^2$ when $a_s$ increases towards $+\infty$.

##### Relative root mean square errors

From previous results, we derive asymptotic predictions on relative root mean squared errors (RRMSE) of estimates, defined as $\mathrm{RRMSE} = \frac{\sqrt{\mathrm{MSE}(\theta)}}{\theta}$ :

- $\mathrm{RRMSE}(a_s)$ increases towards $+\infty$ like $\sqrt{\frac{2}{\mathrm{tr}(R^2)}\frac{a_s}{d_{\min}}e^{\frac{d_{\min}}{a_s}}}$ as $a_s \to 0$;
- $\mathrm{RRMSE}(a_s)$ converges to $\sqrt{\frac{2N}{N-1}}$ as $a_s \to +\infty$;

### 3 Minimum pairwise distance

The sampled area in our study is a equilateral triangle with side length $\sqrt{3}$, which implies an area of $2 \times \frac{1}{2}\frac{\sqrt{3}}{2} \times \frac{3}{2} = \frac{3\sqrt{3}}{4}$. The number of smapling points $N$ is kept constant. Therefore the sampling density of designs is $\lambda = \frac{4N}{3\sqrt{3}}$.

##### Grid design

In a regular grid design, the minimum pairwise distance corresponds $d_{\min}$ to mesh size. A grid design made of equilateral triangles with mesh size $d_{\min}$ reaches density $1/(d_{\min}^2\frac{\sqrt{3}}{4})$ in the plane. Thus, using the expression of density $\lambda$ associated to sampling effort $N$ given above, one obtains that :

$$d_{\min} = \left(\frac{N}{3}\right)^{-\frac{1}{2}}$$

##### Random design

Define $\mathcal{P}(\lambda)$ a Poisson point process with homogeneous density $\lambda$ defined on the plane. Define $D_{\min}(i)$ the minimum distance between a focal point $i$ and the other points of the design. We assume that $D_{\min}(i)$ can be well approximated by the distribution of the analogous random variable computed for a focal point $i$ in $\mathcal{P}(\lambda)$. Then the probability distribution function of $D_{\min}(i)$ should approximately be:

$$\mathbb{P}(D_{\min}(i) < d) \approx 1 - e^{-\pi\lambda d^2}$$

Corresponding density is :

$$f_{D_{\min}(i)}(d) \approx 2\pi\lambda d e^{-\pi\lambda d^2}$$

and approximate expectation is

$$\begin{aligned}\mathbb{E}\left[D_{\min}(i)\right] &\approx \int_0^{+\infty} 2\pi\lambda x^2 e^{-\pi\lambda x^2}\mathrm{d}x \\ &= \pi\lambda\sqrt{2\pi\tfrac{1}{2\pi\lambda}}\int_{-\infty}^{+\infty} x^2 \tfrac{1}{\sqrt{2\pi\frac{1}{2\pi\lambda}}} e^{-\frac{x^2}{2\frac{1}{2\pi\lambda}}}\mathrm{d}x \\ &= \pi\lambda\sqrt{2\pi\tfrac{1}{2\pi\lambda}}\tfrac{1}{2\pi\lambda} \\ &= \tfrac{1}{2\sqrt{\lambda}}\end{aligned}$$

The minimum pairwise distance in the sampling design is $D_{\min} = \min_{i\in\{1,\ldots,N\}} D_{\min}(i)$. Making the approximation that the $D_{\min}(i)$ are independent, one obtains:

$$\mathbb{P}(D_{\min} < d) \approx 1 - e^{-N\pi\lambda d^2}$$

which implies:

$$\mathbb{E}\left[D_{\min}\right] \approx \frac{1}{2\sqrt{N\lambda}} \approx \frac{27^{\frac{1}{4}}}{4N} = \frac{1}{4\times 3^{\frac{1}{4}}}\left(\frac{N}{3}\right)^{-1}$$

$\mathbb{E}\left[D_{\min}\right]$ is therefore always lower than the $d_{\min}$ value obtained in grid design. The relative difference between these two designs increases with sampling effort $N$.

##### Fractal design

A fractal design is obtained by applying $\log(N)/\log(3)$ times the similarities presented in main text. The $k$th application ($k \geq 2$) of similarities yields $d_{\min} = \rho^{k-1}(1-\rho)\sqrt{3}$, therefore:

$$d_{\min} = (1-\rho)\sqrt{3}\left(\frac{N}{3}\right)^{\frac{\log(\rho)}{\log(3)}}$$

Using $\rho = x\sqrt{3}/(2+\sqrt{3})$:

$$d_{\min}(x) = \left(1 - x\sqrt{3}/(2+\sqrt{3})\right)\sqrt{3}\left(\frac{N}{3}\right)^{-\frac{1}{2}-\frac{\log((2+\sqrt{3})/3)}{\log(3)}+\frac{\log(x)}{\log(3)}}$$

Parameter $x$ allows arbitrarily decreasing $d_{\min}$ for given sampling effort $N$. For $x = 1$, one obtains:

$$d_{\min}(1) = \left(2\sqrt{3}/(2+\sqrt{3})\right)\left(\frac{N}{3}\right)^{-\frac{1}{2}-\frac{\log((2+\sqrt{3})/3)}{\log(3)}}$$

which is already smaller than the $d_{\min}$ value of grid. Then $d_{\min}(x)$ decreases toxards 0 as $x$ decreases. Numerically, one obtains that $d_{\min}(x)$ becomes lower than $\mathbb{E}[D_{\min}]$ of random design when $x \approx 0.25$ for $N = 27$.

### 4 Shortest spanning path of fractal and grid designs

Here we derive the length of the shortest spanning path, i.e. a path that covers all sites with minimum distance, for fractal and grid designs. We denote the length of the shortest paths $D^*$

#### Grid design

We consider a grid design covering the triangular sampling area of main text, with $n$ sites on an edge ($n = 7$ in main text). This implies that the total design has $N = n(n+1)/2$ sampling sites. A spanning path is necessarily made of $N-1$ straight lines. The length of each line is by definition above $d_{\min}$, the minimum distance between two sites in the design. Consequently, the length of the shortest spanning path is necessarily larger than $(N-1)d_{\min}$. Reciprocally, in the case of grid design it is easy to find spanning path with a length $(N-1)d_{\min}$. Consequently, we know that the length of shortest spanning paths is exactly $(N-1)d_{\min}$. Using that $d_{\min} = L/(n-1)$, and that $n = (\sqrt{1+8N}-1)/2$, we obtain that:

$$D^*(N,L) = \frac{2(N-1)L}{\sqrt{1+8N}-3}$$

#### Fractal design

We consider a triangular fractal design covering the triangular sampling area with side length $L$ of main text. We call $n$ the number of scales in the design, i.e. the number of iterations of the similarities ($n = 3$ in main text).

Sampling points in the design form equilateral triangles within equilateral triangles within equilateral triangles etc. We call $d_{1,n}$, the side length of smallest equilateral triangles, $d_{2,n}$ the side length of the second smallest triangles, etc. up to $d_{n,n}$, the largest equilateral triange of sampling points. We denote $D_{k,n}$ the smallest distance between two triangles of length side $d_{k,n}$.

Side lengths $d_{k,n}$ follow the recursive relation:

$$\begin{aligned} &d_{0,n} = 0 \\ &d_{k,n} = \rho^{n-k}(1-\rho)L + d_{k-1,n} \end{aligned}$$

which yields the general term:

$$d_{k,n} = L\rho^{n-k}(1-\rho^k)$$

Distance between triangles is:

$$D_{k,n} = d_{k+1,n} - 2d_{k,n} = L\rho^{n-k-1}\left(1 - 2\rho + \rho^{k+1}\right)$$

Below we ask that $D_{k,n} > d_{k,n}$, which is equivalent to:

$$\rho\frac{1-\rho^k}{1-\rho} < \frac{1}{2}$$

which is true for all $k \in \mathbb{N}$ if and only :

$$\rho < \frac{1}{3}$$

which implies, in terms of contraction parameter $x$:

$$x < \frac{2+\sqrt{3}}{3\sqrt{3}} \approx 0.72$$

Under this condition, the shortest patch across the design is necessarily of form : an optimal path within a subtriangle of side length size $d_{n-1,n}$, a move to another subtriangle with length $D_{n-1,n}$, an optimal path within the new sub triangle of side length size $d_{n-1,n}$, a move to the last subtriangle with length $D_{n-1,n}$, an optimal path within the last subtriangle of side length size $d_{n-1,n}$. Calling $D^*(k, n)$ the shortest spanning path wihtin a triangle with side length $d_{k,n}$ in the design, the following recursive relationship holds:

$$D^*(n) = 2D_{n-1,n} + 3D^*(n-1)$$

which yields the general term :

$$D^*(n) = 2\sum_{k=0}^{n-1} 3^k D_{n-k-1,n}$$

Using that

$$D_{n-k-1,n} = L\rho^k\left(1 - 2\rho + \rho^{n-k}\right)$$

one finally obtains :

$$D^*(n) = L\left(2\left(1-2\rho\right)\frac{1-(3\rho)^n}{1-3\rho} + \left((3\rho)^n - \rho^n\right)\right)$$
